# Paradigm of Vanadium pentoxide nanoparticle-induced autophagy and apoptosis in triple-negative breast cancer cells

**DOI:** 10.1101/810200

**Authors:** Parvathy Radhakrishna Pillai Suma, Renjini A. Padmanabhan, Srinivasa Reddy Telukutla, Rishith Ravindran, Anoop Kumar G. Velikkakath, Chaitali D. Dekiwadia, Willi Paul, Sachin J. Shenoy, Malini Laloraya, Srinivasa M. Srinivasula, Sheshanath V. Bhosale, Ramapurath S. Jayasree

## Abstract

Chemo-resistance remains the main hurdle to cancer therapy, challenging the improvement of clinical outcomes in cancer patients. Therefore, exploratory studies to address chemo-resistance through various approaches are highly rewarding. Nanomedicine is a promising recent advancement in this direction. Comprehensive studies to understand the precise molecular interactions of nanomaterials is necessary to validate their specific “nano induced” effects. Here, we illustrate in detail the specific biological interactions of vanadium pentoxide nanoparticles (VnNp) on triple-negative breast cancer cells and provide initial insights towards its potential in breast cancer management at the cellular level. VnNp shows a time-dependent anti-oxidant and pro-oxidant property *in vitro*. These nanoparticles specifically accumulate in the lysosomes and mitochondria, modulate various cellular processes including impaired lysosomal function, mitochondrial damage, and induce autophagy. At more extended periods, VnNp influences cell cycle arrest and inhibits cell migration potentiating the onset of apoptosis. Preliminary *in vivo* studies, on exposing healthy Swiss albino mice to VnNp demonstrated normal blood parameters, organ distribution, and tissue redox balance which further indicated the absence of any adverse organ toxicity. Hence, we foresee tumor-targeting VnNp as a potential drug molecule for future cancer management.

---

Vanadium is a trace element in the human body playing important roles in the normal functioning of the thyroid gland,^1,2^ maintaining calcium homeostasis,^3^ glycemic balance,^1,4^ and helps in fat metabolism^5^. However, the exact role and mechanism of action in the human body are not fully understood. Vanadium enters the body through natural food sources such as mushroom, soybean, black pepper, various seafoods such as shellfish and sea cucumber.^1,6^ Vanadium exists in many forms, like, sodium orthovanadate (Na_3_VO_4_), vanadyl sulphate (VOSO_4_.xH_2_O), sodium metavanadate (NaVO_3_) and vanadium pentoxide (V_2_O_5_).^7^ Vanadium sulfate is found to elicit insulin-mimetic activity^8^ and is also used as a nutritional supplement by athletes for building muscle mass.^8^ Bulk vanadium has proven to have a higher affinity towards adipose tissue^9^ and hence exploited in understanding the possibility of using the same for breast cancer management without the help of any additional targeting. Moreover, it is proven in a case-controlled human study that vanadocene complex treatment in women significantly reduced the chances of breast cancer development^10^. However, the exact mechanism behind the anticancer property of this complex on breast cancer and its overall influence on subcellular organelles, cells, tissues and organisms are yet to elucidate.

Cancer cells rapidly undergo mutations such that they develop resistance to the conventional anti-cancer drug molecules, which emerge as a severe point of concern in cancer management. Developing nanomedicine through the rational design of nanomaterials could address this issue. With the advancements in synthesis and characterization techniques, engineered nanostructures have become popular in a variety of biomedical applications like nanomedicine^11^, bioimaging^12–14^ as well as in various other theranostic applications^15–17^. Some of the previously reported intracellular effects of nanomaterials include the generation of oxidative stress^18^ and induction of autophagy^19,20^. Also, most of the nanomaterials exhibit concentration-dependent structural and functional alterations in various biomolecules like lipids, carbohydrates, proteins, DNA^21^ and even affect cellular organelles. These events modulate the cell death pathway by involving lysosomal dysfunction^22^, mitochondrial damage^23,24^, etc. However, detailed studies on the exact fate of nanomaterials within the cells and its nano-bio interaction studies are still lagging. Due to the lack of such in-depth reviews, nanomedicine has not yet matured from its infancy even after a research history of many decades. Based on this knowledge and existing knowledge gaps, we have investigated the specific activity of vanadium nanoparticles on breast cancer cells within well-controlled biological systems. We hope that an in-depth understanding of the effect of vanadium nanoparticles on breast cancer cells could help to engineer newer generations of nanoformulations, thereby enabling nanoparticle-mediated cancer management.

## Results

### Synthesis and characterization

The synthesis of vanadium nanoparticles followed a hydrothermal route. Subsequently, characterized for understanding its physical, chemical, and morphological properties. Transmission electron microscopy (TEM) revealed the spherical morphology of the synthesized vanadium nanoparticles (VnNp) with an average particle size of 50 nm (Fig. 1a). Energy dispersive x-ray (EDX) analysis confirmed that the sample contained only vanadium and oxygen without any traces of impurities (Fig. 1b). The x-ray powder diffraction (XRD) pattern confirmed the monocrystalline, orthorhombic sherrebanite structure for VnNp following the PDF 00-041-1426 data (Fig. 1c & Supplementary Fig. 1a). Two predominant peaks observed at 20 and 26 degrees represented the preferred orientation of atoms along 001 and 110 planes of the crystal structure. The binding energies obtained from x-ray photoelectron spectroscopy (XPS) at 517 eV and 530 eV correlated to the V2p_3/2_ and O1s electronic orbitals respectively. These binding energies were characteristic of the +5 oxidation state of the sample (Fig.1d & Supplementary Fig. 1b). Raman spectral (RS) analysis showed the typical bending vibrations of O–V=O at 143, 282, and 406 cm^-1^. The stretching vibrations observed at 692 and 995 cm^-1^ denoted doubly coordinated oxygen in V=O and the terminal unshared oxygen atom in V–O respectively (Fig. 1e). Infrared (IR) spectroscopy also revealed specific metal-oxygen stretching and bending modes complementing the Raman data. Peaks observed at 1014, 846 and 671 cm^-1^ represented V=O stretching, O–V vibrations and asymmetric and symmetric stretching modes in V–O–V bridging respectively (Supplementary Fig. 1c). VnNp showed weak absorbance at 430 nm in the UV-visible spectrum (Supplementary Fig. 1d) which is in agreement with earlier reported value^25^ and noted a surface charge of −13 MeV through the zeta potential analysis.

**Figure 1.**
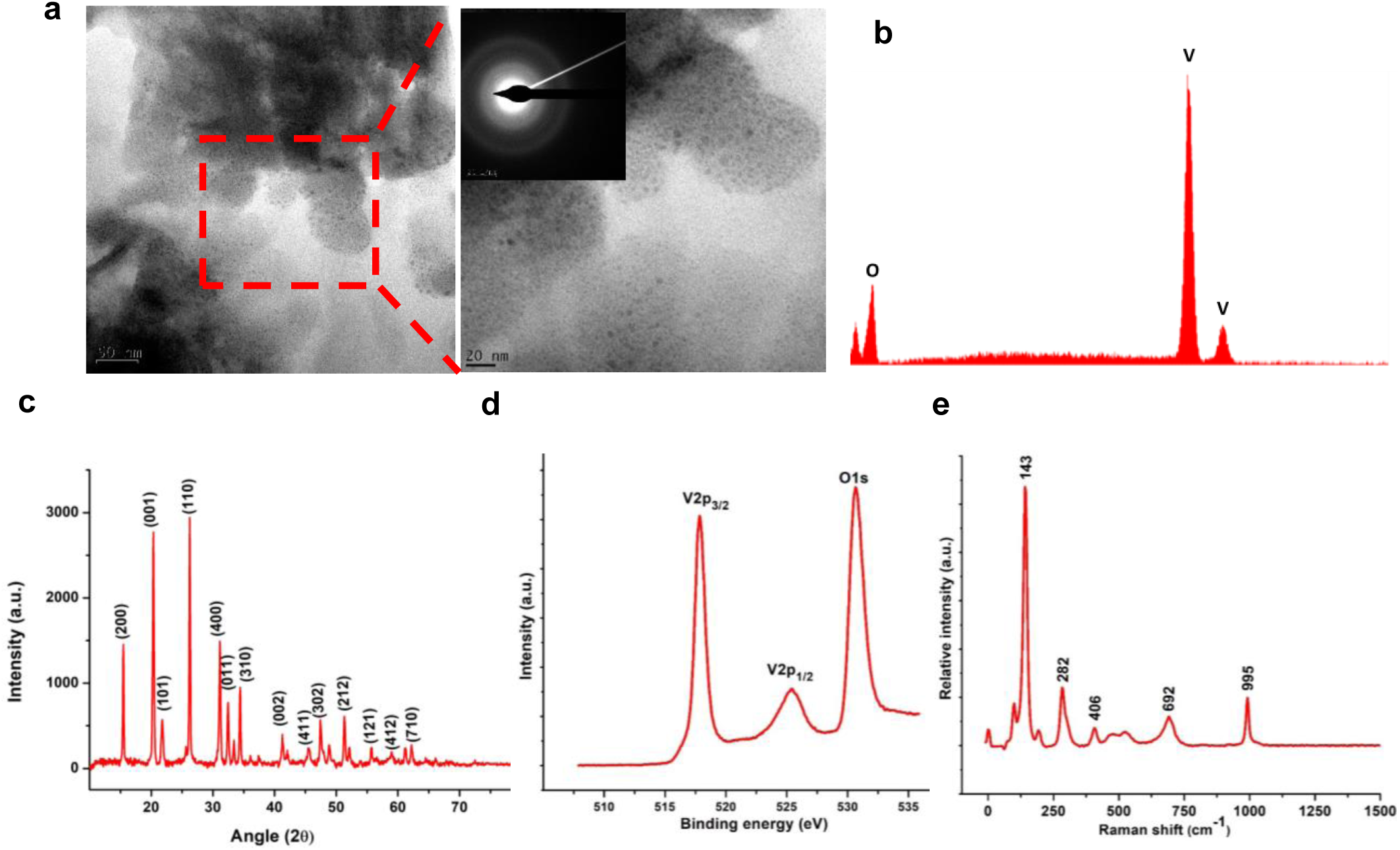
Characterisation of VnNp. (**a**) TEM images depict morphology of the VnNp with an average particle size of 50nm. Smaller particles are shown as zoomed images and the characteristic SAED pattern given in inset. (**b**) Energy dispersive x-ray (EDX) spectra depict the sole peaks corresponding to V and O signifying sample purity. (**c**) XRD pattern demonstrating the orthorhombic crystal structure as per the PDF 00-041-1426 data. (**d**) XPS spectra depict binding energies at 517eV and 530 eV corresponding to V2p_3/2_ and O1s respectively proving the +5 oxidation state. (**e**) Raman spectra showing peaks corresponding to the vanadium oxygen stretching and bending vibrations.

### Dose optimization and cellular internalization of VnNp

A preliminary cytotoxicity assay of VnNp was performed on triple-negative breast cancer cell line MDA-MB-231 and standard fibroblast cell line, L929 by following ISO 10993-5 standard using the 3-(4, 5-dimethylthiazol-2-yl)-2, 5-diphenyltetrazolium bromide (MTT) assay. At a treatment concentration of 50 µM VnNp, the MDA-MB-231 cells resulted in 28.4% and 59.6% cell death towards 24 h and 48 h of treatment. While normal fibroblast cells, L929 showed, only 7.4% and 15.4% cell death at 24 h and 48 h, respectively (Fig. 2a & Supplementary Fig. 2a). Intracellularly sequestered VnNps were observed as distinct small aggregates within the cellular cytoplasm (Fig. 2c) upon 4h of treatment. On to the weak optical property and the high electron density of VnNp, TEM has been used to understand the cellular uptake and cell-material interactions visually. Raman spectral mapping of VnNp treated cells at 4h also demonstrated a characteristic peak of vanadium-oxygen stretching vibrations at 995 cm^-1^ (Fig. 2b) confirming VnNp uptake. Further, flow cytometry analysis of 50 μM VnNp treatment at 4h showed a typical shift in the side scatter pattern (SSC) owing to the increased cellular granularity which could be due to efficient VnNp uptake (Fig. 2d & e). The higher rates of VnNp uptake by cancer cells could be due to the higher levels of metabolic activity of these cells. Meanwhile, the L929 cells failed to show such a change in the SSC area (Supplementary Fig. 2b) where the cells experienced reduced levels of VnNp uptake explaining the observed less cell death pattern in the MTT assay. The drastic difference noted in cell response between healthy and cancer cells for the same concentrations of VnNp intensified the quest for further investigations to prove its suitability for breast cancer management. Based on these results, the interaction of an optimum level of 50 µM or lower concentrations of VnNp on MDA-MB-231 cells were further investigated in detail.

**Figure 2.**
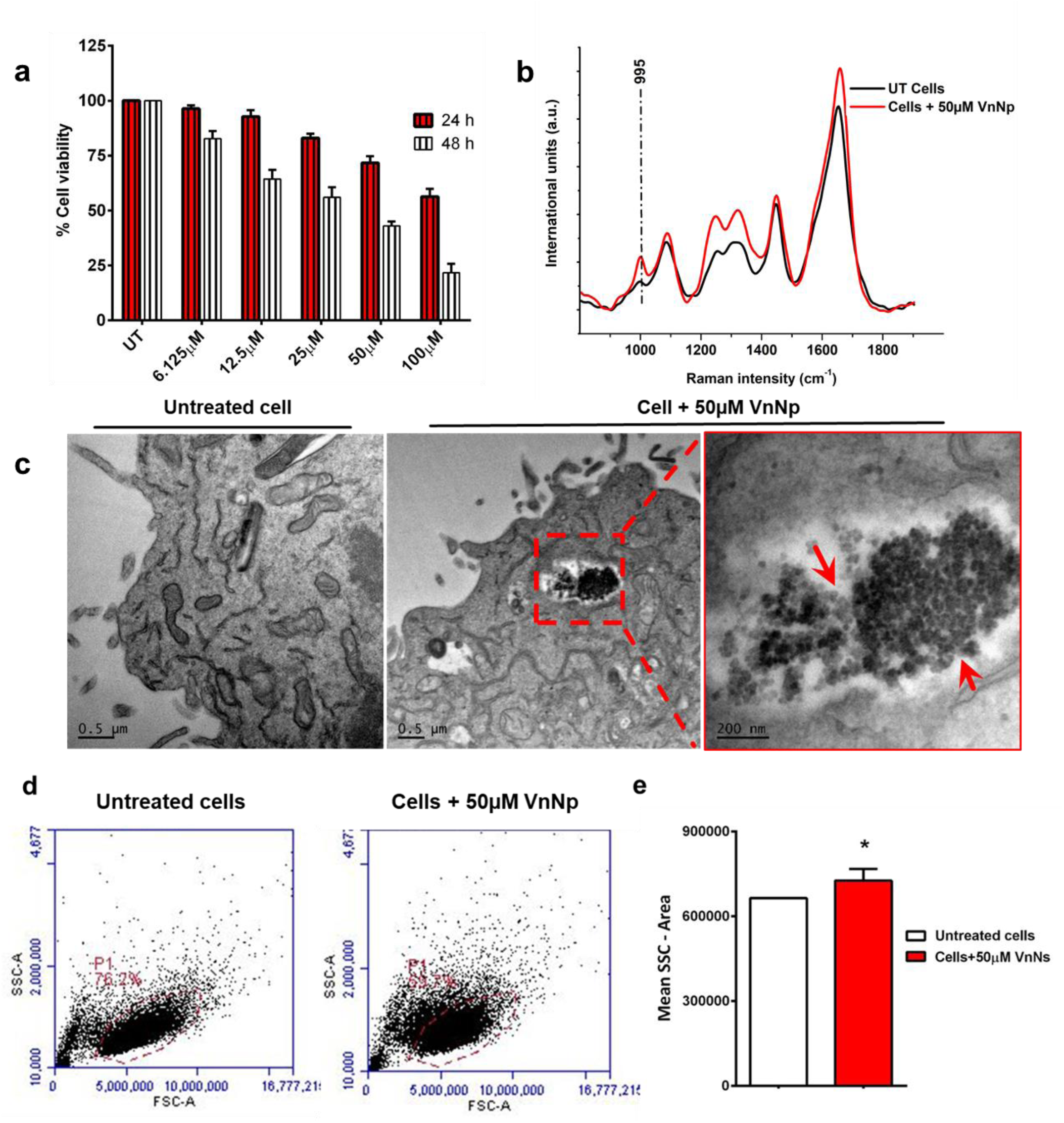
Internalization of VnNp in MDA-MB-231cells. (**a**) MTT assay show percentage cell viability upon 24 and 48h of VnNp treatment. (**b**) Raman spectrum from cells treated with 50 µM VnNp for 4h indicate peak at 995 cm^-1^corresponding to the V-O stretching vibration (**c**) TEM images show untreated cells with intact cellular morphology and organelle architecture and 4h VnNp pre-treated cells with efficient nanomaterial internalization. Zoomed image shows intracellular accumulation of VnNp. (**d**) Scatter plots from the flow cytometry analysis in untreated and VnNp treated cells denote shift in the side scatter pattern indicating VnNp uptake. (**e**) Quantitative representation of cell number from the side scatter region. Statistics: students t-test (*p < 0.01).

### VnNp mediated *in vitro* cellular response

Nanoparticles, once they are in the cytoplasm, often known to stimulate a variety of stress response reactions such as activation of autophagy, ROS generation, etc. These stress responses could result in organelle damage leading to cell death^26^. Autophagy could also be elicited as a cell’s pro-survival mechanism to meet the nutritional requirements of cells at times of nutritional deficiency or other stress cues. This process involves digesting its own damaged organelles such as mitochondria to enable nutrient recycling. During autophagy, autophagosomes are formed with the involvement of an essential protein, the microtubule-associated protein one light chain 3 (LC3) comprising of the free LC-I variant and membrane-bound LC3-II variant. The cells maintain a constant flux between these two forms, by remaining stable and persistent until degradation within the lysosomes. Hence, these marker proteins are used to monitor the extent of autophagy induction in VnNp treated cells. For visual identification of the levels of LC3 protein, stable MDA-MB-231 cells expressing basal levels of Green Fluorescent Protein-LC3 (GFP-LC3) were generated and used. VnNp treatment triggered autophagy induction, where the protein established an active conjugation with the phosphatidylethanolamine on the surface of the autophagosome, forming membrane-bound LC3-II indicated as green fluorescent GFP-LC3 punctae in Fig. 3a. While untreated transfected cells were visualized to have a diffuse green signal throughout the cytoplasm. The extent of autophagy induction was determined based on the expression of the relative number of fluorescent autophagosomes in cells at any time point. The Hank’s Balanced Salt Solution (HBSS) and 25 µM VnNp pre-treated groups did not show visible puncta formation while the number showed an average of 10 punctuates per cell in the 50 µM VnNp treated cell population even from 4h post-treatment, indicating significant autophagy induction (Fig. 3a, b). Consequent to this, the mitochondrial digestion, another hallmark of autophagy was simultaneously checked using tetramethylrhodamine-methyl ester (TMRM). A time-dependent inverse proportionality was observed between the mitochondrial health and LC-3 punctuate formation as indicated by the reduced red fluorescence intensity in the cells treated with VnNp, in the time-lapse images (Fig. 3b).

**Figure 3.**
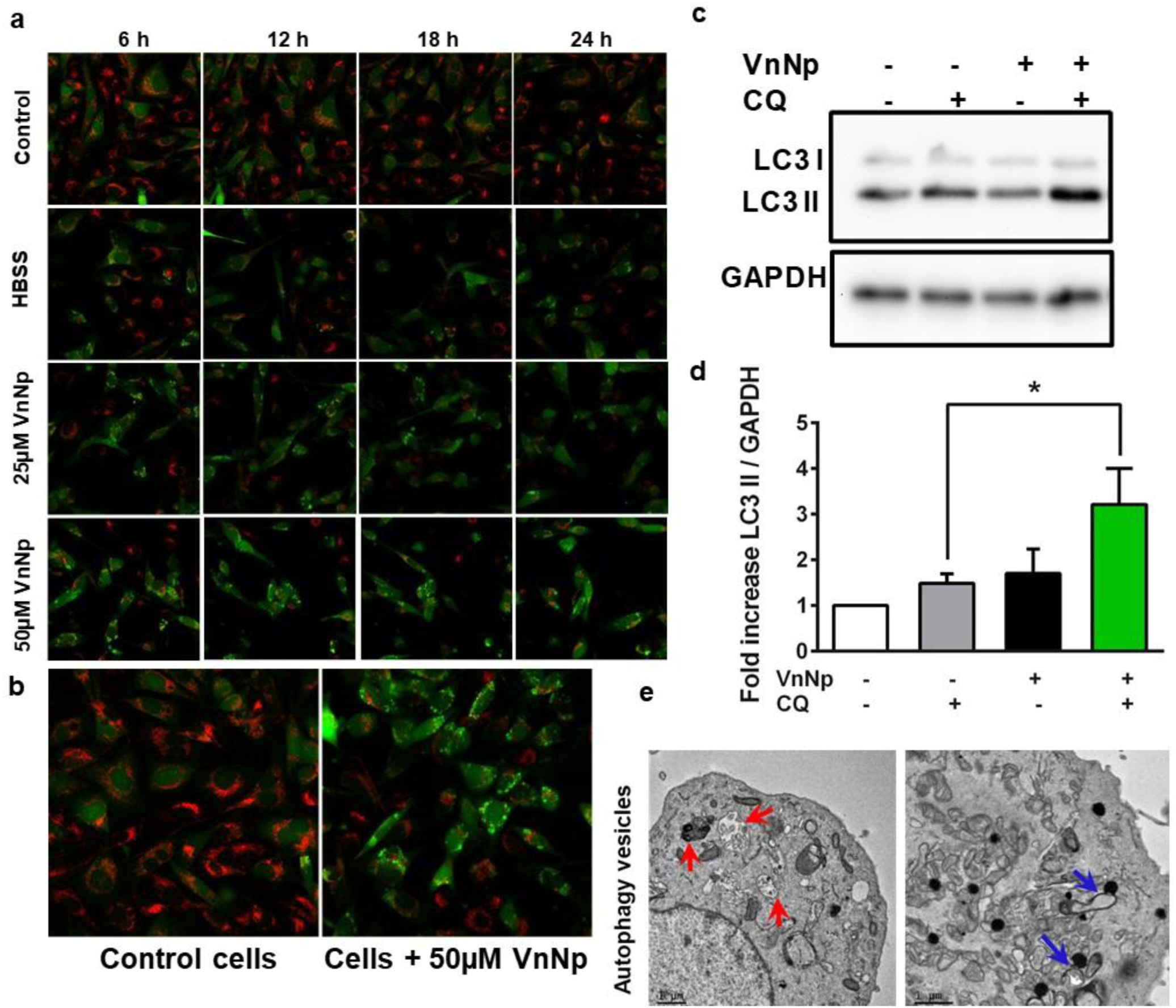
VnNp induced autophagy in MDA-MB-231 cells. (**a**) Time-lapse fluorescence microscopic images on GFP-LC3 transfected cells upon VnNp treatment from 6 to 24h depict the formation of GFP-LC3 punctae and mitochondrial viability indicated from the red signals of indicator dye TMRM. The untreated cells expressed faint, diffuse green signals throughout the cytoplasm alond with strong red fluorescence from healthy mitochondria. While the VnNp treated cells exhibited sharp green spots from autophagosomes and showed a loss of red fluorescence indicating mitochondrial damage upon autophagy induction. (**b**) Depict early LC3 punctae formation in cells with 4h, 50µM VnNp treatment. (**c**) Western blot analysis depict LC3 flux with a prominent increase in LC3-II levels post 50µM VnNp treatment. (**d**) Fold increase in the LC3 expression calculated by normalizing the levels of LC3 II to GAPDH levels, compared the difference in LC3-II levels between samples with and without chloroquine treatment in the flux assay. Data represented is with one way Anova, S.D, n=4, **P < 0.01, *P < 0.05. (**e**) Electron micrographs showing characteristic autolysosome formation and accumulation of cellular organelles (red arrow) and endosome-lysosome fusion (blue arrow), typical indicators of autophagy. Scale bar, 1μm.

Autophagy, a dynamic process was evaluated using a quantitative flux assay, which allowed a comparison between the levels of LC3-I and II proteins at a given time. Western blot analysis also revealed the accumulation pattern of LC3-II protein in response to VnNp treatment further substantiating autophagy induction (Fig. 3c). A lysosomotropic reagent, chloroquine was utilized for the assay which inhibited autophagosome-lysosome fusion by blocking LC3-II degradation, thereby allowing autophagosome accumulation. A two-fold increase in the LC3-II protein levels was observed in the vanadium-chloroquine co-treated cells, indicating autophagy induction by 4h of treatment (Fig. 3d). The expression levels of LC3 protein analyzed until 24 h indicated a visual increase in the LC3-II levels in all the test samples, in comparison with the 24 h untreated and starvation controls (Supplementary Fig. 3). The electron micrographs of VnNp treated cells further identified the presence of cytoplasmic organelles within the autolysosomes (a hallmark of autophagy) which added confirmation of autophagy induction at 4h of treatment (Fig. 3e). Autophagosomes end up fusion with the lysosomes (Fig. 3e) degrading its contents including damaged organelles to meet the energy requirements of the cells. The phago-lysosomal fusion resulted in increased lysosomal size, identified through acid vesicle specific LysoTracker® Red, which further supported autophagy induction in VnNp, exposed cells.

The control cells expressed scattered basal level of lysosomes in fluorescent and electron micrographs (Fig.4 a & b), while the 24 h treated cells showed the formation of gigantic phagolysosomes which labeled very strong with the LysoTracker dye and appeared as large persistent red fluorescent spots even at 48 h VnNp treated cells (Fig. 4c). These acidic vesicles were found to exhibit a pattern accumulation near the peri-nuclear region to maintain the high acidity of organelle associated with VnNp treatment as evident from the electron micrographs (Fig. 4d). A 10 fold increase in the fluorescence intensity was observed in the treated cells compared to the control cell population (Fig. 4e), which indicated increased lysosomal acidification. Here, the occurrence of large acidic vesicles could be due to incomplete lysosomal degradation or defective lysosomal recycling. Increased lysosomal size could elevate its surface tension leading to membrane disruption and subsequent release of acidic contents into the cytosol. However, it was interesting to note that this has not happened, as no acidic membrane disruption was observed even after 48 h of treatment (Fig. 4c). To confirm further, the lysosomal membrane integrity was analyzed using acridine orange (AO), a highly selective metachromatic fluorochrome. Sharp spots of red fluorescence were observed from the VnNp treated cells throughout the study period until 48 h of incubation, indicating intact lysosomal membrane architecture (Supplementary Fig. 4). Together, these findings suggest that even though VnNp treatment led to phagolysosomal accumulation and increased acidic vesicles, they did not elicit lysosomal membrane disruption. However, such a condition could lead to the onset of various cellular stress response reactions in treated cells such as ROS generation. Under normal physiological conditions, ROS in cells is tightly regulated by the strong interplay between various mitochondrial enzymes involved in redox signaling pathways. These proteins facilitate normal homeostasis and protect cells from oxidative stress-induced damage. However, ROS inducers overpass this protective machinery, damage mitochondrial membrane potential, and induce oxidative stress. Hence, the mitochondrial membrane potential (ΔΨ_m_) was quantitatively evaluated using flow cytometry with a fluorescent probe, JC10. A positive control, carbonyl cyanide 3-chlorophenylhydrazone (CCCP) was used as a potential inhibitor of mitochondrial membrane for gating the cells, whereas untreated cells served as negative controls in flow cytometry. The VnNp treated cells demonstrated a substantial increase in the percentage of green fluorescence and a shift in the mean fluorescence intensity from red to green, over a treatment period of 24 h (Fig. 5a) indicating altered mitochondrial membrane potential. Fluorescence microscopy also revealed strong spots of red fluorescence from the intact mitochondria within untreated control cells, and a drop in the red fluorescence intensity pattern with a subsequent increase in the green fluorescence, upon 24 h VnNp treatment (Fig. 5b) which indicate loss of mitochondrial membrane potential. Such membrane damages could further open mitochondrial permeability transition pores (MPTP) and allow more VnNp to accumulate inside the mitochondria. This was further confirmed with the electron micrographs indicating the accumulation of VnNp within the mitochondria of treated cells (Fig. 5c). The distorted mitochondrial membrane could further influence the escape of electrons from the electron transport chain producing highly reactive oxygen radicals like superoxides, peroxides, hydroxyl radicals, singlet oxygen, etc. These radicals once formed rapidly react with the membrane structures and result in loss of physiological activity. Hence, an overall idea on the Oxidative Stress index in VnNp treated cells were tested spectrofluorimetrically using 2’,7’ dichlorofluorescein diacetate (DCFDA) assay. The cells in the VnNp treated group showed approximately 30% decrease in ROS levels at an initial 4h incubation period in comparison with their untreated control population. However, the ROS pattern reversed with a significant increase in green fluorescence towards 24 h treatment in comparison with the corresponding untreated controls (Fig. 5d & e). VnNp mediated reduction in superoxide levels in cells at initial time points was quite unexpected, and hence mechanism behind this was investigated in the cell-free system. Accordingly, the rate of inhibition of pyrogallol auto-oxidation in the presence of VnNp was checked and found a 7% reduction within the initial 10 minutes of incubation (Supplementary Fig. 5). This is attributed to the ability of VnNp to inhibit superoxide formation or accelerated neutralization of oxy-radicals. Subsequently, a much more sensitive Amplex^®^ Red assay was executed to check specifically the levels of H_2_O_2_, in MDA-MB-231 cells to confirm the efficacy of VnNp to act as antioxidants or pro-oxidants against intrinsic levels of ROS. This assay also clearly indicated an initial decline in the amplex red fluorescence intensity pattern which further established that VnNp is responsible for the initial decline in the H_2_O_2_ levels *in vitro* (Fig. 5f). Nonetheless, the levels of H_2_O_2_ elevated towards 24 h treatment in a similar pattern as that was observed in the DCF assay.

**Figure 4.**
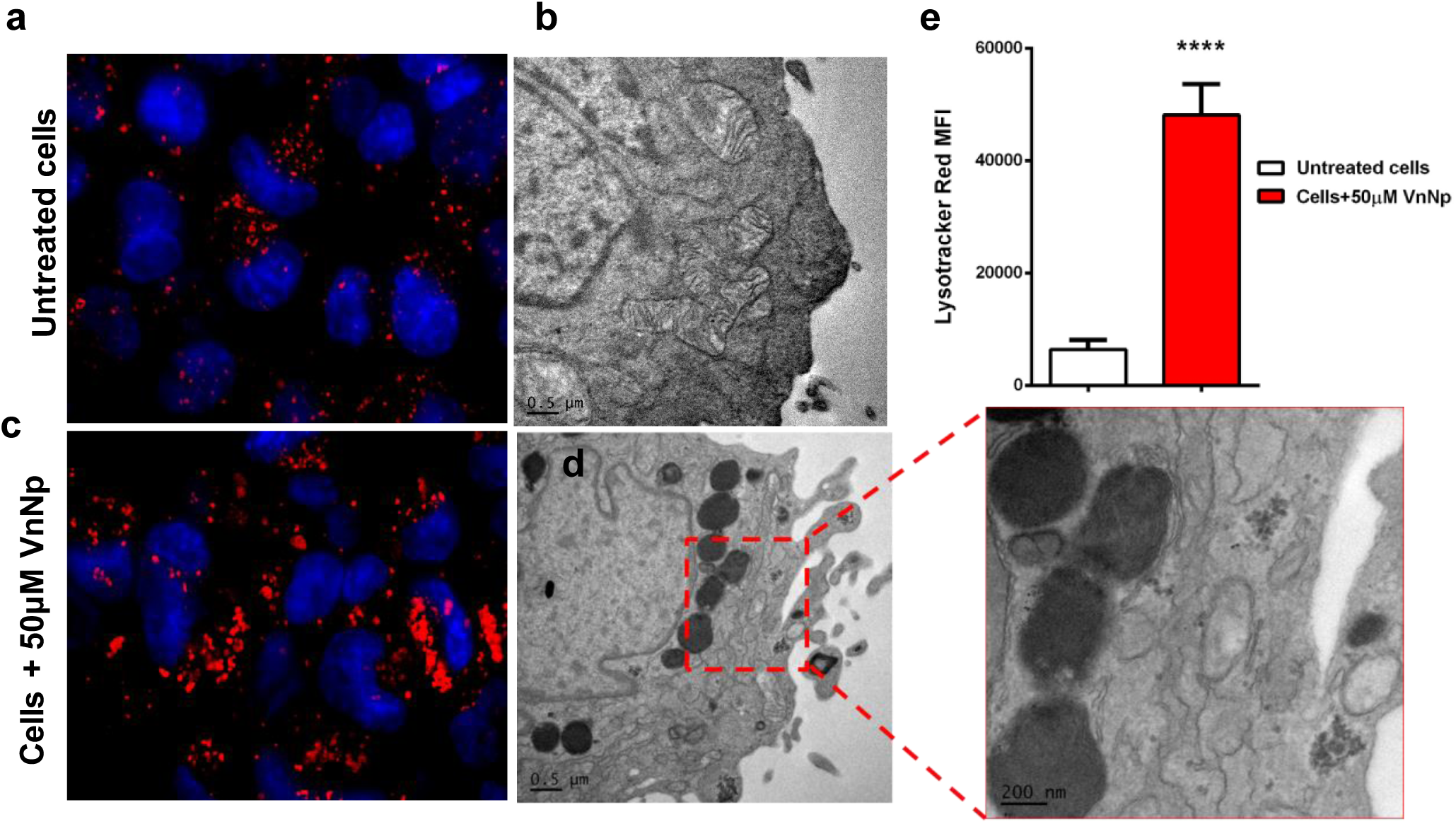
Lysosomal behaviour upon VnNp treatment. (**a & b**) Fluorescent microscopic image and electron micrograph showing untreated MDA-MB-231 cells showing normal lysosomal number and size. (**c & d**) Fluorescent microscopic image and electron micrograph showing changes in the lysosomal organisation upon 48h of 50μM VnNp treatment. Indicate a strong difference in lysosomal size, total lysosomal number and increased red fluoresce intensity pattern in fluoresce image and also from electron micrographs as shown in the zoomed image. (**e**) Students t-test showing statistical significance between mean fluorescence intensity patterns of untreated controls with that of the 50μM treated cells. Data represented from independent experiments considering a minimum of 50 different focal points, error bar shows S.D where n=6, ****P < 0.0001.

**Figure 5.**
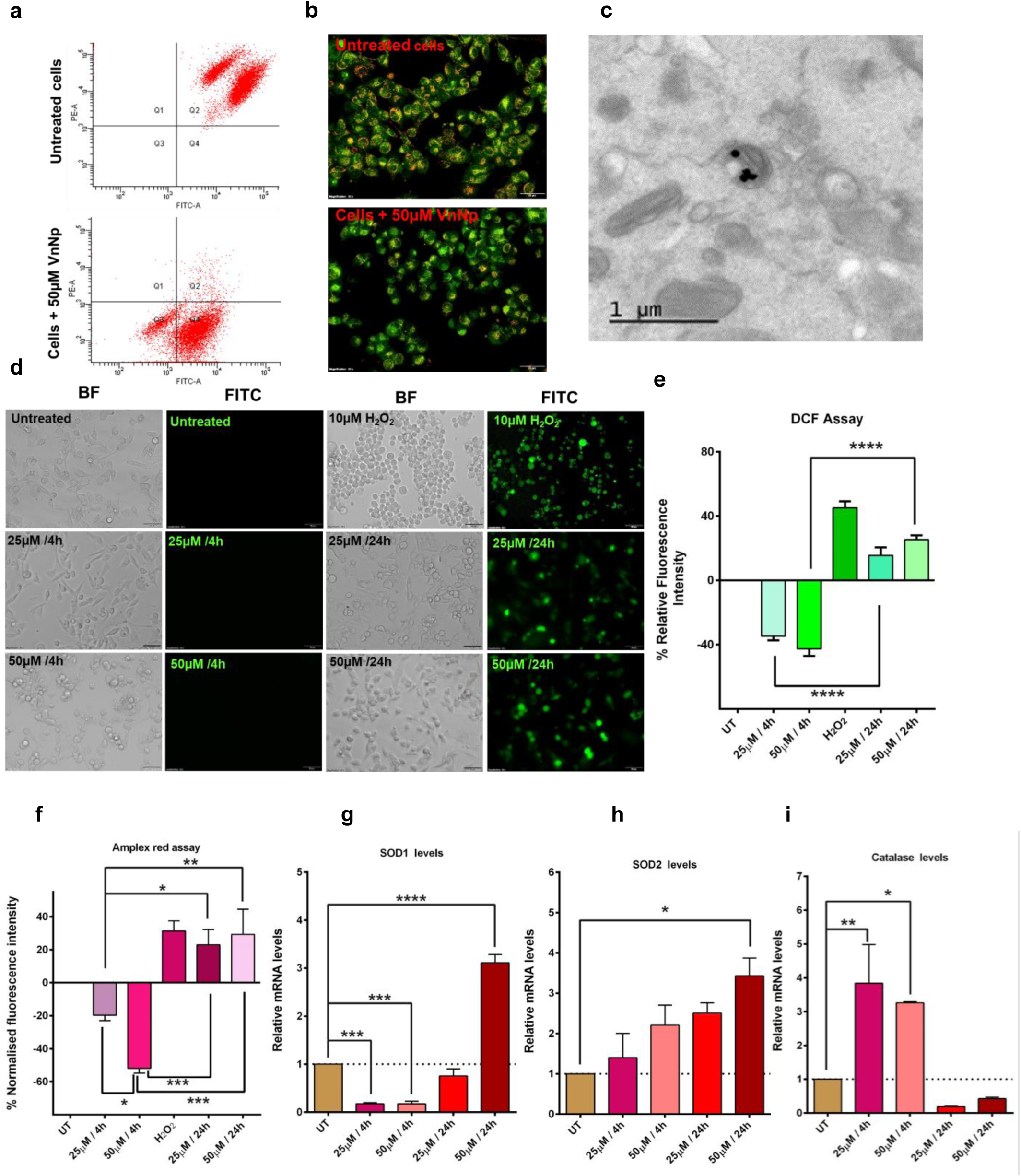
Influence of VnNp on mitochondrial redox balance. (**a**) Flow cytometry analysis indicate altered mitochondrial membrane potential (ΔΨm) in the JC-10 assay where healthy untreated cells were seen towards both red (TRITC) and green (FITC) channels. While the VnNp treated cells were seen more towards the green channel indicating loss of mitochondrial membrane potential. (**b**) Fluorescence micrographs depict sharp spots of red fluorescence along with diffuse green signals from healthy cells (top panel) and VnNp treated cells showed loss of mitochondrial membrane potential, indicated through the less intense red fluorescence from cells in the bottom panel in JC-10 stained cells. (**c**) Electron micrograph depict the presence of VnNp inside mitochondria of treated cells. (**d**) Fluorescence microscopic images of DCFH-DA staining for ROS detection in H2O2 treated and VnNp treated cells. Scale bar 10µm. (**e**) Spectrofluorometric quantitative measure of green fluorescence with DCF-DA show the percentage cells with varying levels of ROS. (**f**) Amplex red assay shows quantitative measure on the specific levels of H_2_O_2_ at 4 h and 24 h of VnNp treatment, cells treated with 10µM H_2_O_2_ acted as a positive control. (**g**) Show SOD1 mRNA levels towards 4h and 24h of VnNp treatment, indicate an increase in mRNA levels only towards 24h. (**h**)) mRNA levels of SOD2 shows a gradual time dependent increase in levels towards 24h. (**i**) catalase mRNA levels shows an initial increase and a gradual drop within 24h of incubation time. All the data were analysed using two way Anova, data representing error bar shows S.D, where n=4, ****P < 0.0001, ***P < 0.001, **P < 0.01 & *P < 0.05.

The catalytic property of vanadium pentoxide was reported to vary with the planar arrangement of vanadium and oxygen^27^. Moreover, the catalytic activity and the antioxidative cell response corresponding to the 010 plane of nanowires have been in reports^25^. Accordingly, it was established that the 010 plane features a haloperoxidase activity, where it acts as a Lewis base capable of accepting oxygen atoms and the 001 plane stacked with neutral coordinatively saturated vanadium cations^27^. Hence, the observed cellular response of the present study is expected to attribute significantly to the planar arrangement of atoms, size, shape and extent of nanomaterial uptake at any given point of time. The present study showcases the ROS quenching activity of VnNp during the first hours of treatment which could be attributed to the preferred orientation of the 110 plane of the crystal structure as seen in the XRD pattern (Fig. 1c). This plane has cleaved V-O covalent bonds, and resultant unsaturated V and O atoms at its side surface make them more easily hydroxylated and contributes to the initial ROS quenching activity^27^. However, the quenching activity reduces towards 24h, which could be due to the loss of surface reactivity pertaining to surface saturation. Additionally, prolonged uptake of VnNp could have activated the cellular defense machinery, thereby elevating intracellular ROS levels.

The intrinsic cellular antioxidative response was also accounted for by studying the mRNA levels of various SOD’s and catalase as they act as indirect indicators on the presence of oxy-radicals and H_2_O_2_. Nanomaterial interaction with the cellular system includes a complex dynamic process that impairs the enzymatic balance between various dismutases like SOD1, SOD2, SOD3, and catalase. Where, SODs are involved in the conversion of reactive oxygen to comparatively less responsive H_2_O_2_ and catalase facilitates the transformation of H_2_O_2_ into the water and molecular oxygen. Elevated level of ROS is characteristic of cancer cells, hence the pattern of stress response mRNA levels upon VnNp treatment were monitored in MDA-MB-231cells using real-time PCR (RT-PCR) analysis. Relatively low levels of SOD1 mRNA was observed in cells treated with 25 & 50 μM VnNp for 4h and 25μM for 24 h, while a two-fold increase in mRNA levels was observed with 50μM for 24 h treatment (Fig. 5g). A regular pattern of growth in the mitochondrial SOD2 was also observed from 4h until 24 h of post-treatment (Fig. 5h). This observation is possibly attributed to the initial basal level quenching activity of ROS upon VnNp treatment which could subsequently account for the low initial levels of SOD mRNA. A decrease in surface reactivity of VnNp could have increased ROS levels leading to higher levels of SOD mRNA towards 24 h of treatment (Fig. 5g). Also, relative levels of catalase mRNA revealed more than 2-fold increase in its levels with 25 & 50 μM of VnNp treatment at 4h. However, the catalase mRNA levels dropped towards 24 h post-treatment (Fig. 5h). Here, an initial surge in the catalase mRNA reiterates our finding of decreased H_2_O_2_ from the corresponding amplex red signals within 4h of treatment (Fig. 5f & 5h). Altogether, the results indicate an initial drop in the ROS levels attributed to the superoxide dismutase like activity of VnNp pertaining to its structure. Besides, VnNp activates initial intrinsic levels of catalase thereby resulting in decreased H_2_O_2_ levels towards the first hours of treatment, all working in tandem.

These elevated levels of ROS and cellular responses could cause direct or indirect damage to the DNA exhibiting transient cell cycle arrest. Hence, implications of cellular stress response in cell cycle progression were ascertained with flow cytometry analysis using 50 μM or lower concentrations of VnNp. Any DNA damage or incomplete chromatin replication in the late S phase was quantified by analyzing the DNA content at various time points. Interestingly, a dose-dependent progressive increase in cell number towards the G2/M phase of the cell cycle was observed with 50 and 25 μM of VnNp treatment for 24, 36 and 48 hours in comparison to their respective untreated control population (p < 0.05) (Fig. 6a). The cell number in the G1 phase declined from 66% (control) to 43% (50 μM) in the treatment group (Fig. 6b) and a corresponding increase in the G2/M phase from 15% to 38% post 24 h treatment. A similar pattern of cell cycle arrest was also observed at 36 and 48 h, which indicated time-dependent irreversible cell cycle arrest at the G2/M phase post 24, 36 and 48 h of VnNp treatment.

**Figure 6.**
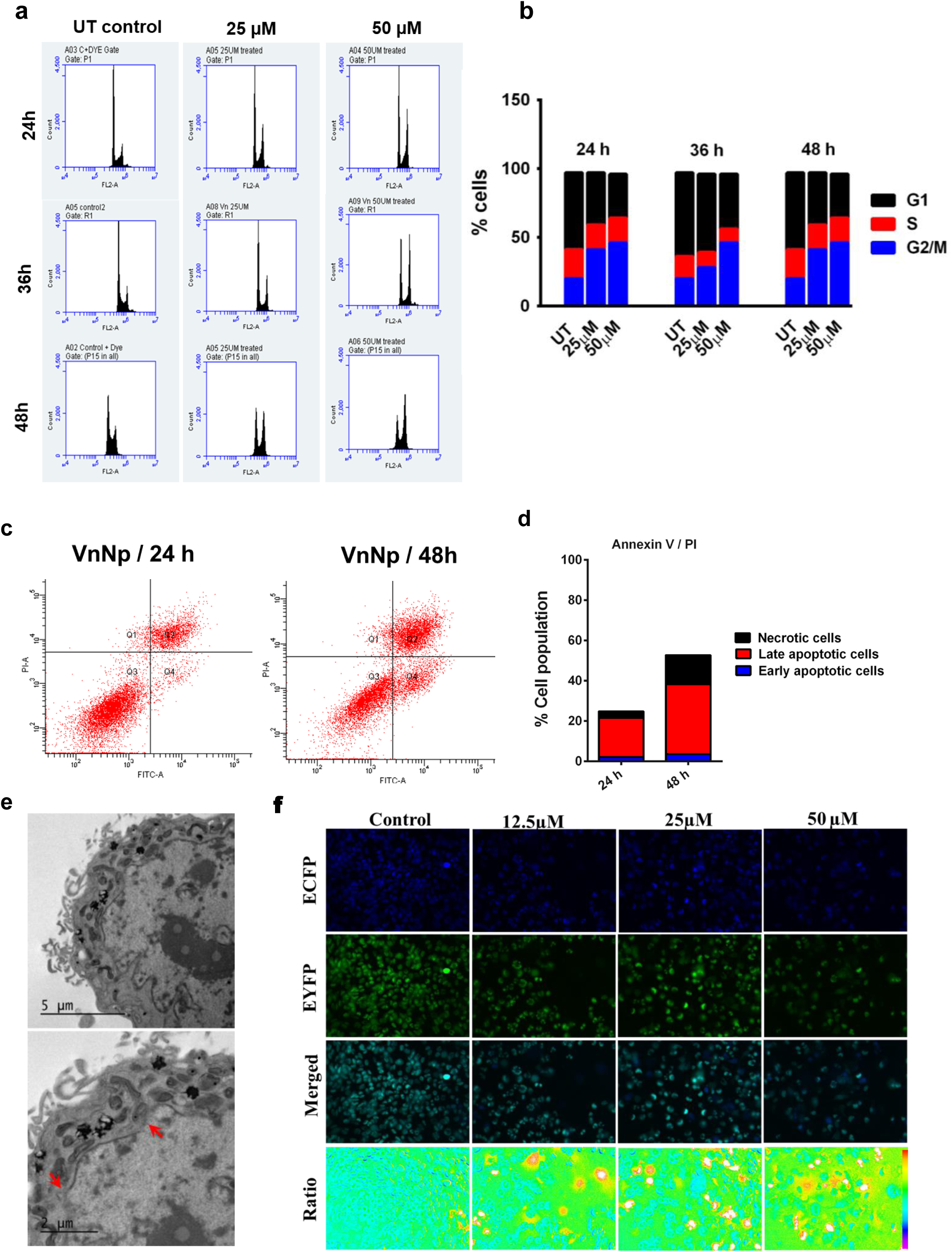
VnNp mediated cell cycle arrest and apoptosis induction. (**a**) Show the flow cytometry data on cells at different phases of cycle in cells treated with 25 & 50μM of VnNp for 24, 36 & 48h. Indicate cell cycle arrest with an increase in cell population in the G2/M phase in each of the histograms represented. (**b**) Data on quantification from the flow analysis represent the number of cells at each phase of cell cycle. (**c**) Flow cytometry analysis indicate the number of cells at different phases of cell death through annexin-V/PI assay. (**d**) The quantitative data from the flow analysis shown represents cells treated with VnNp for 24h and 48h with percentage cells in the early apoptosis, late apoptosis and necrosis. (**e**) Electron micrographs depict dissolution of nuclear membrane in cells treated with 50μM VnNp for 48h. (**f**) A FRET based caspase activity assay on cells stably expressing SCAT3 NLS. Ratiometric images indicate significant increase in the number of caspase activated cells upon treatment with 50μM VnNp for 48 h. Imaged using a 20X objective with 0.75 NA.

Together, with these results on cell cycle arrest and increased ROS levels in treated cells, we anticipated that the cell becomes no longer capable of effectively repairing the cellular damage and end up in the cell death pathway. The percentage of cells at different stages of cell death was quantified using Annexin-V-FITC / Propidium Iodide (PI) dual staining assay with flow cytometry. The number of cells in the late apoptotic phase increased from 19.4% to 34.7% as the VnNp treatment progressed from 24 h to 48 h (Fig. 6c & 6d). Also, the electron micrographs depicted leaky nuclear membrane morphology in cells that are exposed to 50 μM of VnNp for 48 h (Fig. 6e) indicating apoptotic induction. Moreover, the extent of cytosolic caspase activity in apoptosis triggered cells was monitored using a FRET-based live-cell imaging technique which is insensitive to the environmental changes. The assay used MDA-MB-231 cells engineered with a caspase-3 sensor FRET probe, SCAT3 NLS. Cells pretreated with VnNp induced caspase activation and subsequent FRET loss, which is reflected from the increased donor (ECFP) to acceptor (Venus) fluorescence ratio in Fig. 6f and Supplementary Fig. 6. Here, these cells exhibited a characteristic FRET loss and an increase in ECFP/Venus ratio, whereas intact cells exhibited FRET activity towards 48 h of treatment. Studies on understanding the cytoskeletal architecture and nuclear condensation pattern revealed actin filament reorganization and chromatin condensation with 50 μM VnNp treatment for 48 h (Fig. 7a & Supplementary Fig. 7 & 8). Various apoptotic cellular events were also marked through membrane blebbing (Fig. 7b & Supplementary Fig. 8) nuclear membrane disruption (Fig. 6e) and nuclear fragmentation (Supplementary Fig. 9) all supportive of the apoptotic mode of cell death towards 48h of VnNp treatment.

**Figure 7.**
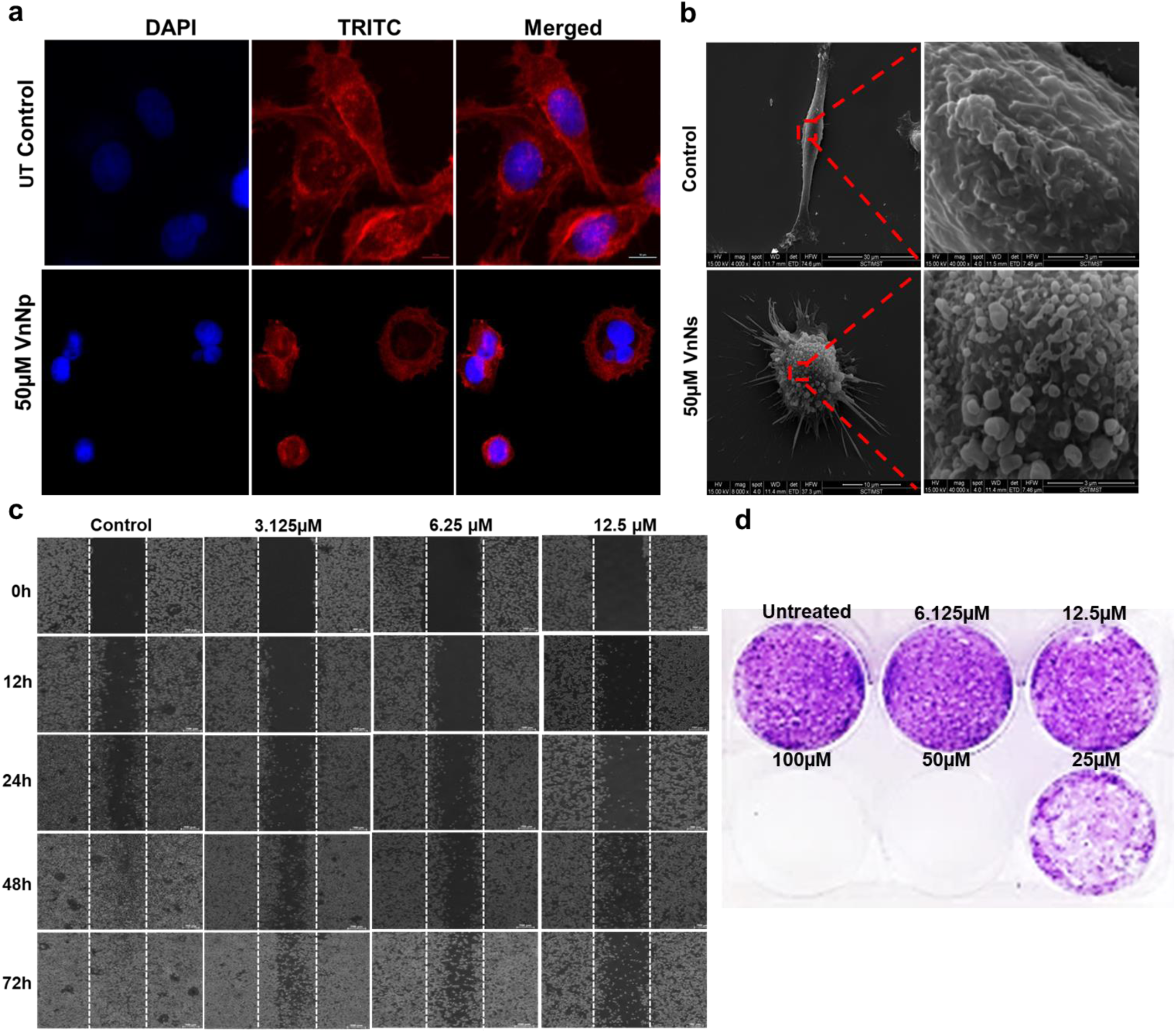
Prolonged exposure to VnNp on cells. (**a**) Fluorescent micrographs depict actin filament reorganization and chromatin condensation in 50μM VnNp treated cells in comparison with the untreated controls observed at 48h of post treatment. (**b**) Representative SEM micrographs show surface morphology, characteristic loss of cell morphology and membrane blebs indicating apoptotsis induction in VnNp treated cells for 48h, n=3. (**c**) Bright field images on cells in the wound healing assay. Indicate inhibition of cell migration with 3.125, 6.25 and 12.5 μM of VnNp treatment for various time points. (**d**) Clonogenic assay depict inhibition on cell colony formation in 50 and 100 μM VnNp treatment and a reduction in cell clone intensity with 6.25, 12.5 and 25 μM of VnNp concentrations.

**Figure 8.**
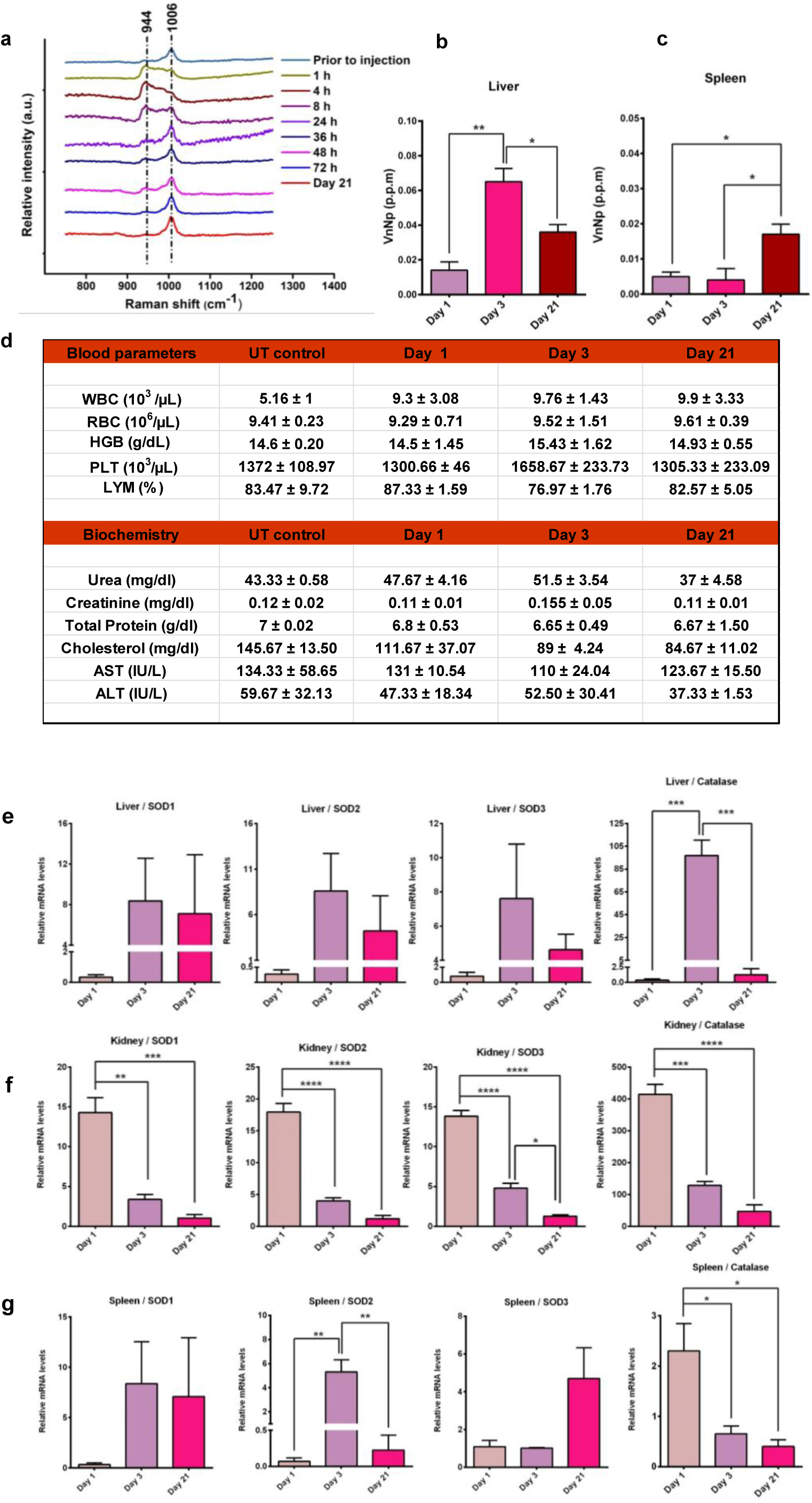
Evaluation on the *in vivo* influence of VnNp. (**a**) Raman spectra show prominent urea peak in urine sample collected from animals prior to VnNp injection and a peak shift was observed in samples after VnNp treatment collected at different time points. (**b**) ICP analysis indicating the presence of traces of elemental vanadium in liver at various time points (**c**) ICP analysis indicating the presence of traces of elemental vanadium in spleen at various time points (**d**) Blood analysis on histology and biochemical parameters between samples from UT controls and treated groups. (**e**) RT-PCR analysis on relative mRNA levels of SOD’s and catalase in the liver samples towards day 1, 3 and 21. (**f**) RT-PCR analysis on relative mRNA levels of SOD’s and catalase in the kidney samples towards day 1, 3 and 21. (**g**) RT-PCR analysis on relative mRNA levels of SOD’s and catalase in spleen samples, indicating normal levels by the end of 21-day of post exposure. Error bar indicates S.D, where n=5, ****P < 0.0001, ***P < 0.001, **P < 0.01 & *P < 0.05.

Through this study, a systematic mechanistic insight is given on how triple-negative breast cancer cells behave towards the VnNp treatment with an initial attempt of unaccomplished cell survival, finally resulting in cell death. We also tried to understand whether VnNp inhibits breast cancer cell migration *in vitro* through a wound-healing assay in MDA-MB-231 cells maintained in minimal FBS to prevent cell proliferation. Interestingly, it was observed that the wound did not patch even after 72h of treatment with 12.5, 6.25 and 3.125 µM VnNp. Conversely, the control group covered up the patch through enhanced cell migration towards the wounded area within 72 h (Fig. 7c). Hence, we are optimistic that VnNp could inhibit breast cancer cell metastasis, which is yet another critical area of concern to be addressed in its management. A colonogenic assay also exhibited the potential of VnNp to inhibit colony formation (Fig. 7d) which further proved its potential to inhibit tumor growth *in vitro*.

### VnNp mediated *in vivo* response

Nanoparticles largely influence the external environmental conditions to which they have been exposed. Hence, we tried to understand the fundamental biological interactions of VnNp in a multicellular level involving complicated detoxification and excretory mechanisms that are otherwise absent in the *in vitro* scenario. For which, female Swiss albino mice intravenously administrated (*i*.*v*.) with a single dose (1mg/kg) of VnNp were utilized. The extent of renal elimination of VnNp was accounted through Raman spectral analysis of the urine samples collected at various time points after VnNp exposure. The prominent symmetrical C-N stretching bond of urea at 1006 cm^-1^ in the pre-treatment urine samples and metal-oxygen vibration of VnNp at 995 cm^-1^ in the post-treated samples were considered at various time points starting from 1 h of injection. The interaction between VnNp and urea resulted in the peak shift of V-O stretch from 995 to 944 cm^-1^. The peak corresponding to V-O stretch gradually increased until 72 h of VnNp treatment while that of urea gradually decreased. Post three days, the urea peak ultimately regained its position whereas the very weak V-O peak at 72h indicated VnNp clearance, by this time. There was no presence of VnNp in the urine by the 21^st^ day (Fig. 8a). The possible chances of accumulation of VnNp in the excretory organs were further evaluated using elemental analysis with inductive coupled plasma optical emission spectroscopy (ICP-OES). A quantitative measure on the residual levels of vanadium in the vital organs like liver, kidney and spleen was evaluated. The liver samples indicated 0.014 parts per million (ppm), 0.065 ppm and 0.036 ppm of vanadium in the 1, 3 and 21-day post-treatment samples (Fig. 8b). The levels were maximum in the day-3 sample indicating the possibility of its persistence in the blood until deposition and the levels declined towards day-21 which attribute towards particle elimination. The spleen samples showed traces of vanadium, with 0.017 ppm towards day-21 (Fig. 8c) while none of the kidney samples showed detectable levels of vanadium at any of the periods. In addition, blood analysis studies revealed elevated WBC levels in the treatment group in comparison with the untreated control population, which could be attributed to the presence of traces of vanadium in the spleen. Various other hematology and biochemical parameters like creatinine, urea, aspartate aminotransferase (AST) & alanine aminotransferase (ALT) levels were comparable to that of controls in the 21-day samples (Fig. 8d). This reflected normal blood parameters including liver and kidney function marker proteins between the treated and untreated groups. Very interestingly, a statistically significant decrease in cholesterol levels was observed in the 21-day sample in comparison with the control group which could be correlated to the anti-lipidemic activity of vanadium, known from ancient times. As this point needs further investigations and is not under the scope of the present study, no further investigations were done in this regard.

Like the *in vitro* studies on the real-time levels of stress response mRNA, the pattern of SOD’s and catalase from liver, kidney, and spleen were also analyzed to monitor the extent of nanomaterial induced stress responses, *in vivo*. The transient mRNA levels of these genes specify the redox status of organs involved in the elimination process. The low levels of SOD1, 2, 3 and catalase mRNA in the day-1 liver samples indicated no immediate stress response in the liver (Fig. 8e). An increase in levels of all the 3 SOD’s and catalase mRNA was observed towards day 3, indicating delayed response towards VnNp treatment. Interestingly, it was found that all these mRNA levels dropped back to normal by day-21 in liver samples (Fig. 8e). This observation points out that despite the presence of traces of VnNp within the liver even after 21 days following a single dose exposure, the organs still maintained a healthy redox balance, hepatic and renal function. The kidney samples showed a rise in the mRNA levels of all the three SOD’s and catalase in the day-1 sample which was found subsided indicating the absence of any delayed stress response in the kidneys (Fig. 8f) and thus proved the renal safety aspects of VnNp. Even though in kidney samples, an initial increase in SOD mRNAs was observed, the process was also dealt with enormous catalase levels involved in detoxifying any H_2_O_2_ produced during the first-day post-treatment. The relative levels of SOD’s and catalase in spleen samples also indicated basal redox mRNA levels by the end of 21-day of post-exposure (Fig. 8g) indicating the absence of VnNp induced stress response in the spleen. In summary, it can be deduced from these results that the presence of VnNp in none of the excretory organs elicited any stress response. These results were again confirmed with the histopathology observation (Supplementary Fig. 9). The tissue sections obtained from the untreated and VnNp treated samples showed similar tissue organization in the liver and kidney, while the 21-day spleen samples showed elevated levels of lymphocyte infiltration, which could be a reason for high levels of WBC observed in the blood studies.

In conclusion, this study gives a greater understanding of the ultrastructural interactions of VnNp *in vitro* and the mechanistic effects leading to breast cancer cell death. Preliminary insight into the safety and *in vivo* biodistribution aspects are also demonstrated. A series of cellular events like alterations in the cytoplasm, specific organelles, and nuclei of the cells after VnNp treatment are explained. VnNp activates autophagy response causing mitochondrial damage and late apoptosis without lysosomal membrane disruption. The treatment also modulates the redox milieu by affecting SOD and catalase mRNA levels and result in efficient neutralization of H_2_O_2_. In addition, VnNp efficiently blocked cell migration and tumor growth *in vitro*. Further, the *in vivo* murine experimental analysis with VnNp administration revealed normal mRNA levels of stress response genes in the liver and spleen. Also, the treatment neither caused tissue damage nor affected their function even after three weeks of a single shot tail vein injection. Thus, the study provides a mechanistic understanding of the interaction of VnNp *in vitro* and *in vivo*, as a platform for the development and design of vanadium-based nanotherapeutics for breast cancer management, thereby strengthening the need for detailed material interaction studies for biomedical applications. The study is an attempt towards developing a promising nano-vanadium-based anticancer agent, and can be modified for future application as a promising anticancer molecule with extensive studies on tumor-bearing animal models to validate their potential to be used in nanomedicine.

## Materials and Methods

### Synthesis and characterization of VnNp

Synthesis followed a hydrothermal method utilizing Vanadium oxychloride and benzyl alcohol in the ratio 1:40, a reaction set at 120°C, refluxed for 4 days. A thick precipitate obtained, washed in absolute ethanol and ultrapure water following overnight drying. The product ground into a fine powder and calcined at 450°C to yield the final product as a yellow powder. The material was thoroughly characterized for crystallinity from the Powder X-ray diffraction pattern. XRD on a D8 Advance model machine (Bruker AXS, Germany) using Cu Kα radiation at the tube parameters of 40 kV and 30 mA, at a scan speed of 4 degrees/min. The binding energies of vanadium and oxygen in vanadium pentoxide analyzed with XPS analysis using a MULTILAB 2000 Base system with X-Ray, Auger, and ISS attachments, Thermo Scientific. SEM and EDX spectra recorded with FEI-Quanta 200. High-resolution TEM images and selected area electron diffraction pattern was recorded with JEOL 2100 operating at 200kV. Infra-Red Spectra recorded using transmission mode and KBr pellet method in Cary 600 (Agilent Technologies). Raman spectra from the bare VnNp as well as vanadium pretreated cells were analyzed using a confocal Raman Microscope (alpha 300R, WITec Inc. Germany) with the spectral Imaging Mode. Raman mapping of cells was performed with cells grown over tin-coated glass plates, treated with the nanomaterial.

### Cell culture and maintenance

MDA-MB-231 and L929 cells were kind gifts from RGCB, Trivandrum. Cultured and maintained in DMEM medium (GIBCO-Invitrogen, NY) supplemented with 10% fetal bovine serum (PAN-Biotech GmbH, #3302) with penicillin G (100 mg/mL), and streptomycin (100 mg/mL) in a humidified CO_2_ (5%) chamber at 37 °C. Cell culture medium was replaced every two days, and upon 80% confluence, cells were subcultured with 0.25% trypsin-ethylenediaminetetraacetic acid (trypsin-EDTA) (Invitrogen) or were further used for cell studies.

### Cell viability assay

MTT assay evaluated the percentage of cell viability with various concentrations of VnNp. Cells were seeded at an initial seeding density of 1 × 10^4^ cells per well in a 96 well culture plate and maintained for 24h at 37°C and 5% CO_2_ (ambient conditions). VnNp dispersed at a concentration of 1mg/ml in phosphate-buffered saline pH at 7.4 was used as a stock solution of 0.5mM. The cells without any treatment served as the control and 100μL of a particular concentration of VnNp in media served as the treatment group. Maintained for 24/48h under similar conditions and then followed standard instructions from Invitrogen. Used a microtiter plate reader (Synergy H1 hybrid multi-mode microplate reader, Bio-Tek) operating at 570 nm and a reference wavelength of 630 nm. Results were normalized to untreated control values, and the percentage relative cell viability was calculated.

### Transmission electron microscopy (TEM)

All solvents and chemicals for cell fixation were purchased from ProSciTech, Australia unless otherwise mentioned. Cells treated with VnNp were pelleted by centrifugation at 1500 rpm and transferred to sterile PCR tubes. And followed 0.1 M cacodylate buffer washes (X3) and fixed in 2% paraformaldehyde with 0.1M cacodylate buffer (pH 7.4) and 2.5% glutaraldehyde for 2 h. Post-fixation, in 1% osmium tetroxide in distilled water for 1.5 hours at room temperature, cells were washed and dehydrated in a series of ethanol concentrations (from 50 to 100%) for 10-minute intervals and finally in absolute acetone for 30 minutes. Infiltrated through a series of increasing levels of acetone: Spurr’s resin overnight and eventually infiltrated twice through 100% fresh Spurr’s resin and allowed to polymerize in an oven pre-set at 70 °C for 48h to obtain resin-embedded cell blocks. The blocks were sectioned using a diamond knife fitted to an ultra-microtome (Leica) into 70-90nm nm thick sections and transferred into copper grids. Electron microscopy was performed using JEOL-JEM 1010 TEM operating at a voltage of 80 kV with magnifications from 15-30,000x.

### Expression vectors and generation of stable cells

SCAT3-NLS vector was a kind gift from Dr. Masayuki Miura, University of Tokyo, Japan. pEGFP-LC3 was a gift from Tamotsu Yoshimori (Addgene plasmid # 21073). The cell transfection was performed with Lipofectamine LTX (Invitrogen, #15338-100) as per the manufacturer’s protocol. Cells stably expressing the transgene were selected by antibiotic G418-Geneticin® (Invitrogen, #11811-031) selection in 500 µg/ml for three to four weeks. Cells with homogeneous transgene expression were sorted and expanded using BD FACS Aria III cell sorter (BD Biosciences).

### Imaging EGFP-LC3 and TMRM

Cells stably expressing EGFP-LC3 were grown in 96-well optical bottom culture plates and stained with 100 nM TMRM prior to VnNp treatment. VnNp at appropriate concentrations were prepared in 5% FBS containing phenol-red free DMEM supplemented with 20 nM TMRM. HBSS supplemented with 20 nM TMRM served as a positive control. For imaging, cells were sequentially scanned using a laser scanning Leica TCS SP8 WLL Confocal microscope (Leica Microsystems, Wetzlar, Germany) equipped with supercontinuum white-light laser. A 488 nm and 561 nm excitation lines from white light were used for imaging EGFP-LC3 and TMRM respectively. The emission at 500-550 nm was collected while exciting at 488 nm using GaAsP detectors. For TMRM, the emission at 600 ± 25 nm was collected while exciting at 561 nm. Cells maintained at 37°C, optimum humidity and 5% CO_2_ in a microscope incubation chamber (Leica Microsystems, Wetzlar, Germany).

### Protein extraction and western blot analysis

VnNp pretreated MDA-MB-231 cells were harvested in ice-cold PBS and pelletized at 6500rpm for 3 min at 4 °C. Lysed in lysis buffer (50mM Tris-Cl, 150mM NaCl, 1mM EDTA, 1% (v/v) Triton X-100), 1mM Phenyl Methane Sulfonyl Fluoride (PMSF P7626) and protease inhibitor cocktail (Sigma P8340) incubated in ice for 30min followed by 10 min centrifugation at 10,000rpm, 4°C, collected supernatants, determined concentrations through the BCA Protein Assay kit (Thermo Scientific 23225). Loaded equal amounts of boiled, cooled proteins (30 μg) in Laemmli buffer onto 12% SDS-polyacrylamide gel and transferred to polyvinylidene difluoride (PVDF) membrane as per standard protocol. Primary antibodies diluted 1:1000 and secondary antibodies 1:5000 for western blotting (WB). Probed with rabbit primary antibody, LC3B (Cell Signalling Technology 2775s) overnight at 4 °C, washed (3X) with TBST. Subsequently incubated with HRP conjugated Goat anti-rabbit secondary antibody (Invitrogen 65-6120) for 1hr at room temperature, and washed in TBST (3X). GAPDH anti-mouse antibody raised in Rabbit (Abgenex 10-10011) and secondary antibody (Invitrogen 61-6520) served as the loading control. The bands developed using ECL reagent (Millipore WBKLS0500) observed under chemiluminescence in a BIO-RAD ChemiDoc XRS, quantified using Quantity One software.

### Lysosomal staining and fluorescence microscopy

Cells were seeded onto sterile coverslips at an initial seeding density of 1×10^5^ cells per well and allowed to grow overnight. Treated with VnNp at appropriate concentrations and stained with lysosome specific LysoTracker Red™ dye (Invitrogen, USA). The cells without any nanomaterial treatment served as the untreated controls. The average maximum intensity from each field was obtained from the z-stack of 8 slices by selecting average mean fluorescence intensity from at least 10 different regions of interest (ROI) at a constant laser exposure of 300 ms on both control and treatment groups. Imaging was carried out using 540-550nm BP excitation and 575-625nm BP emission filter with a 570nm dichromatic mirror using an Olympus Fluorescence Microscope (IX83, Tokyo, Japan) equipped with a metal halide lamp (X-Cite, series 120PC Q) and a cooled CCD camera (XM10, monochrome, Olympus). Image analysis was performed using CellSens Imaging software (Olympus).

### Mitochondrial membrane potential assay

For flow cytometry, cells were seeded into 6 well plates at an initial seeding density of 1 × 10^6^ cells/ml. Treated with specific concentrations of VnNp and subsequently trypsinized, recovered and resuspended in 500 μL of 1X JC-10 dye-loading solution kept at room temperature and allowed to incubate in the dark for 20 minutes. The dye forms aggregate within intact mitochondria and emit red fluorescence while it remains as monomers in the cytoplasm and give out green fluorescence from cells with altered mitochondrial membrane potential. The cells were analyzed immediately using a BD FACS Aria III (BD Biosciences) with FL1 channel for green fluorescence and FL2 channel for orange to red fluorescence. Qualitative red fluorescence intensity from treated cells were visualized using microscopy images were also obtained using personalized blue excitation (470-495nm), red emission (575-625nm) BP filters and 570nm dichromatic mirror equipped in an Olympus Fluorescence Microscope (IX83, Tokyo, Japan) having a metal halide lamp (X-Cite, series 120PC Q) and a cooled CCD camera (XM10, monochrome, Olympus).

### Measure of oxidative balance

The rate of inhibition on Pyrogallol auto-oxidation in the presence of VnNp was performed through kinetic studies with UV-1800 Shimadzu UV-Vis spectrophotometer in a cell-free system. The protocol was adopted from the improved pyrogallol assay experiment.

Reactive oxygen species (ROS) in cells subsequent to VnNp exposure were measured using H_2_DCFDA reagent (Invitrogen cat# C6827). Cells were seeded on a 96 well culture plate at an initial seeding density of 2×10^4^ cells/well and allowed to grow overnight. The VnNp treatment followed a similar treatment concentration as that of MTT assay and 10 μM H_2_O_2_ treatment for 20 minutes served as positive controls. The assay was performed as per the manufacturer’s instructions. The ROS levels from untreated cells maintained under ambient conditions were taken into account in order to normalize treatment values and readings from wells with all the said reagents and vanadium except cells were obtained to minimize the background interference and to normalize the readings. The fluorescence intensity was analyzed in a 96-well spectrofluorimeter plate reader (Synergy H1 hybrid multi-mode microplate reader, Bio-Tek) at an excitation and emission wavelength of 492 and 527 nm respectively. Qualitative cellular fluorescence images captured with an Olympus Fluorescence Microscope (IX83, Tokyo, Japan).

The specific levels of H_2_O_2_ produced in cells treated with VnNp were monitored using an Amplex assay red kit from Invitrogen and followed the manufacturer’s instructions.

### Cell cycle analysis

Cells plated at an initial seeding density of 1×10^6^ cells per well in 6 well plates and allowed to attach overnight. The cells treated with 25 and 50μM of VnNp and were subsequently incubated for 24, 36 and 48 h. Recovered the cells following centrifugation at 1000rpm for 5min so as to obtain a cell pellet. Fixed in ethanol and stained with propidium iodide (PI) staining buffer containing 200mg PI (Sigma), 0.1% (v/v) Triton X-100, 2 mg DNAse-free RNAse-A (Sigma) in 10 mL PBS for 15 min at room temperature. Further, analyzed immediately for measuring variations in DNA content from the fluorescence of PI. Flow analysis counted 10,000 events on a BD Accuri™ C6 flow cytometer.

### Assay for cellular apoptosis

Cells were seeded at an initial seeding density of 1 × 10^6^ cells /mL per well onto six-well plates. Treated with VnNp with the proper concentrations and maintained for 24 and 48h. Cells were recovered, and the percentage cells at different stages of cell death were quantified using Invitrogen™ eBioscience™ Annexin V Apoptosis Detection Kit following manufacturer’s instructions. Analyzed with BD FACS Aria III (BD Biosciences) excited with a 488 nm laser, where FITC detected in the FL-1 channel with a 525/30 BP filter while PI detected in the FL-2 channel by a 575/30 BP filter. Standard compensation using single-stained and unstained cells and 10,000 events were counted on each set of experiment. Data analysis was done using FlowJo software (Tree Star Inc.).

### Cytoskeletal and nuclear morphology

Cells were cultured over sterile coverslips (Blue star 13 mm) and were treated with VnNp for 48h, fixed with 3.7 % formaldehyde for 15 min and permeabilized (100% methanol, 10 min, −20 °C), washed (PBS, ×3) and blocked (10% BSA in PBS, 30 min, room temperature). Subsequently incubated with Rhodamine-phalloidin (cat. numb: R415) for F-actin and with 5 µg/ml Hoechst dye 33242 (Invitrogen, USA) for 15 min at room temperature for nuclear staining and followed steps as per manufacturer’s instructions. The nuclear and cytoskeletal changes during apoptotic cell death were observed under an N-STORM super-resolution confocal microscope. Cells, fixed and dehydrated in ethanol gradient were further dried and sputter-coated with gold to visualize the morphological changes under SEM FEI-Quanta 200.

### High-throughput screening for caspase activation

Cells expressing SCAT3-NLS were seeded in 96 well optical bottom culture plates (BD Biosciences, #353219) maintained at ambient conditions. They were treated with working concentrations of VnNp in phenol-red free complete medium and imaged with BD Pathway™ 435 Bioimager (Becton Dickinson Biosciences, USA). The excitation/dichroic mirror/emission filter combinations used for imaging were 438±24/458LP/483±32, 438±24/458LP/542±27, for CFP, CFP-YFP FRET respectively. Imaged using a 20X objective with 0.75 NA.

### *In vitro* migration assay

Initial seeding density of 5 × 10^5^ cells per well in 30mm dishes was allowed to attain confluence for 24 h. Gently scraped from one end to end having a constant width. Plates were washed to remove floating cells, treated with different concentrations of VnNp and kept undisturbed in low serum conditions to slow down cell proliferation and maintained for various time points. Further, cell migration across the inflicted wound area was imaged with a phase-contrast microscope (Nikon) at five or more randomly selected fields.

### Clonogenic assay on MDA-MB-231 cells

The anti-proliferative action of VnNp further confirmed through a clonogenic assay. This assay replicates the capability of a single cell to rise into a colony and considered as the standard for determining long-term cell viability as this assay reflects all modes of cell death. MDA-MB-231 cells at the exponential phase were plated at single-cell density (500 cells/well) into 6 well plates and allowed to adhere for 24 h. Following VnNp treatment, cells maintained for 24 h, and the medium replaced with fresh culture medium and cells incubated for further 14 days. Cells washed in PBS, fixed in 4% paraformaldehyde and stained with 0.5% methylene blue in PBS for 30 min and washed several times with distilled water to remove excess dye and the dried plates, photographed with a digital camera.

### Animal housing

Groomed female Swiss albino mice (25±3 g) in individually ventilated cages with six animals per experimental group. Animals were acclimated and allowed to synchronize their estrous cycle 4 weeks prior to the start of experiments. The management of animals was done in compliance with regulations of the Committee for the Purpose of Control and Supervision of Experiments on Animals (CPCSEA). The animal experiments were approved by the Institutional Animal Ethics Committee (IAEC) for animal Care Govt. of India. Approval No: IAEC approval No: SCT/IAEC 243/August/2017/94. Groups randomly divided as vehicle control (saline), and VnNp in saline-injected groups were maintained for 24h, 3 days and 21-day post-injection and euthanized by cervical dislocation. Tissue samples immediately stored in Trizol reagent for RNA isolation and a portion for ICP analysis.

### Urine and blood analysis

The presence of vanadium in urine samples was analyzed using a Confocal Raman microscope Raman (Witec Inc. Germany, alpha 300R) supported with a laser beam focused on the sample through a 60X water immersion objective (Nikon, NA = 1.0) and a Peltier cooled CCD detector. The excitation wavelength of 532 nm achieved with the second harmonic of Nd: YAG laser and spectra collected from 400-4000 cm^-1^ with 1 cm^-1^ spectral resolution. Measurements were done on samples placed over Tin coated glass slides and used WITec Project plus (v 2.1) software package for data analysis. Compared samples from the same mice before and after VnNp administration (n=6).

Whole blood from orbital sinuses of each animal was analyzed for various hematological and biochemical parameters. The assays carried out in a fully automated Beckman coulter system.

### ICP-OES

Analyzed for the presence of vanadium in the liver and kidney at different time points post-injection. Briefly, samples (100 mg) were pre-digested in 5.0 ml concentrated nitric acid (HNO3, ICP grade, Sigma) followed by complete digestion in dilute 2% HNO3 with heating at 150°C. Vanadium (ICP standard 1000ppm Sigma) was used as the internal test standard. Both standard and test solutions were measured using three different wavelengths using ICP-OES (Perkin Elmer 5300DV, USA).

### Quantitative Real-Time PCR

Standard protocols were followed for the quantitative real-time PCR experiments. In short, total RNA was isolated using TRIzol (Invitrogen) reagent, quantified using Nanodrop, and around 2µg RNA was converted into complementary DNA (cDNA) using Superscript VILO First-Strand cDNA synthesis kit (Thermo Fisher Scientific) and qPCR amplification was performed with Power SYBR^®^ Green PCR Master Mix (Life Technologies, CA). Oligonucleotides were designed using NCBI pick primers and were purchased from Ocimum Biosolutions. The relative expression was calculated using ΔΔCT method with 18S rRNA as the endogenous control. The run was performed on a 7900 HT Fast Real-Time PCR system (Applied Biosystems, CA) under standard cycling conditions. Melting curve analysis was done to ensure the product specificity. The primers used in the experiment are listed in Supplementary Table 1.

### Histopathology analysis

The histopathology evaluation was done on the liver, spleen, and kidney collected from each treatment and control group. The tissue samples obtained from mice post euthanasia washed several times in normal saline and immediately transferred into 10% neutral buffered formaldehyde (10% NBF) to prevent tissue degradation. The NBF processed samples cut into small pieces, kept in tissue cassettes and dehydrated in a series of isopropanol and xylene. Subsequently impregnated in molten paraffin and allowed to cool in order to obtain tissue embedded paraffin blocks. Thin tissue sections of 5μM thickness microtome (Leica RM 2125 RT) drifted on warm water picked on clean l-lysine pre-coated microscopic glass slides. The sections stained in Hematoxylin, partially removed by acid-alcohol, counterstained with Eosin dye (H&E) as per standard protocol. Slides mounted in DPX to provide better optical transparency observed under a light microscope (Olympus CX31) with Q-capture ProTM 6 software.

## Supplementary Information

islinked to the online version of the paper at www.nature.com/nature

## Note

The authors declare no competing interests.

## Acknowledgements

P.R.S., thank the Indian Council for Medical Research, Govt. of India for the Junior Research Fellowship (JRF) No.3/1/3/JRF-2012/HRD-148 (20037) and the Commonwealth of Australia for the Endeavour Research Fellowship-2015. J.R.S. & P.R.S. thank the Director, SCTIMST for the infrastructural facilities. S.V.B. acknowledges the Goa University for the award of a Professorship Position under UGC-FRP. The authors thank the Director, RGCB, and Director, IISER Trivandrum for the infrastructural facilities. R.A.P. thank funding from the DST, Govt. of India. R.R. thank the DST Inspire, Govt. of India for JRF. A.K.G.V. acknowledges the post-doctoral research funding from IISER Trivandrum, Govt. of India. P.R.S. thank Dr. T. R. Santhoshkumar and Shankar N.V. of Cancer Research Program, RGCB for their support in getting the GFP-LC3 time-lapse fluorescence microscopic images and caspase activation experiments [Figure 3(a &b) & 6f]. The authors thank Dr. Masayuki Miura, University of Tokyo, Japan, for gifting the expression vector SCAT3-NLS and Tamotsu Yoshimori for pEGFP-LC3 (Addgene plasmid # 21073). P.R.S and S.R.T thank Dr. Ravi Shukla of RMIT for the cell culture facility. The authors thank Mr. Nishad K V for his inputs and technical support towards the SEM analysis and Mr. Sanal Das (Olympus DSS Imagetech Pvt Ltd.) for the technical support and inputs during fluorescence cell imaging. The authors thank Dr. Manoj Komath for the ICP facility, Dr. Suresh Babu and Dr. Remya K.R for their support during ICP analysis. Also, thanks to Dr. Sabareeshwaran for the histopathology analysis. The authors thank Mr. Manoj and Mr. Sunil for their technical assistance in handling and housing of investigational mice. The authors acknowledge the facilities, the scientific and technical support of the RMIT University’s Microscopy & Microanalysis Facility (RMMF), a linked laboratory of the Microscopy Australia for the EM analysis. The authors also recognize the generous support from the RMIT University’s Micro Nano Research Facility (MNRF) for the flow cytometry and confocal imaging facilities.

## Author Contributions

R.S.J., S.V.B., and P. R.S. conceived the idea of the study and designed the work. P.R.S. is involved in the synthesis and chemical characterization of VnNp. R.S.J., S.V.B., and P.R.S interpreted chemical characterization of VnNp and its properties [Figure 1(a-e)]. P.R.S. contributed to images shown in Figures 2a, 2(d-e), 3(a &b), 4(a-e), 5(a-e), 6(c&d), 6f, 7(b&d), 8(b-d). P.R.S., R.R., and A.K.G.V. performed the autophagy assay, evaluated the results under the supervision of S.M.S and contributed to Figures 3(c & d); P.R.S. and S.R.T. performed flow cytometry and confocal imaging as shown in Figure 6(a&b) and 7(a&c). P.R.S. and C.D.D performed electron microscopic analysis on cells and contributed to images shown as Figures 2c, 5c & 6e. P.R.S. and W.P performed Raman spectral mapping shown in Figure 1e, 2b, & 8a. P.R.S. and S.J.S. carried out the *in vivo* VnNp injection and monitored the experiments. P.R.S. and R.A.P. performed amplex red assay and real-time PCR experiments, evaluated the RT-PCR experimental data and formulated the results shown in Figure 5 (f-i) and Figure 8 (e-g) under the supervision of M.L. The manuscript was written with contributions from all the authors and all have approved the final version of the manuscript.

