## Supplementary information for "Paradigm of Vanadium pentoxide nanoparticle-induced autophagy and apoptosis in triple-negative breast cancer cells"

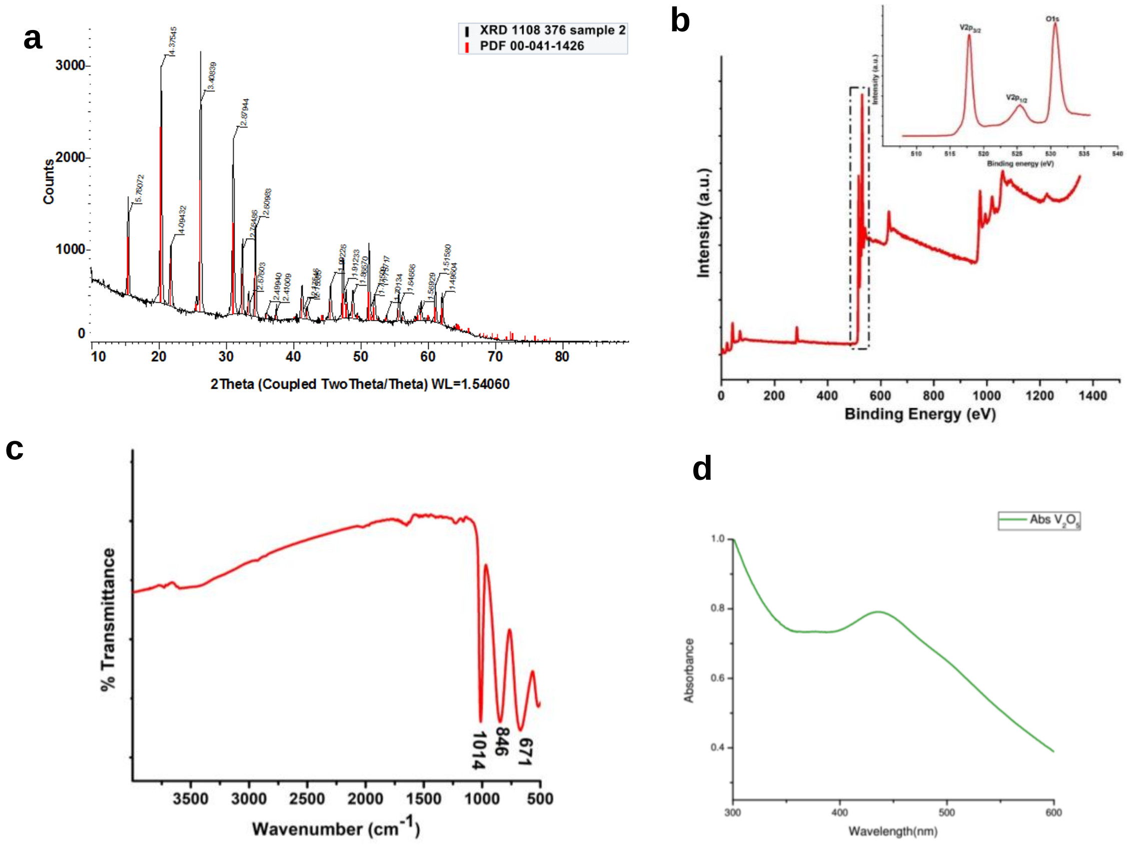


**Supplementary Figure 1. Characterization of VnNs.** (**a**) XRD pattern cross match with the PDF 00-041-1426 data. (**b**) XPS pattern elucidating the orbital arrangement of atoms in the vanadium pentoxide nanomaterial. The figure shows survey spectra with inset depicting high resolution V2p and O1s peak analysis showing the only presence of vanadium and oxygen in the sample indicating high sample purity. (**c**) IR spectra showing characteristic metal-oxygen stretching and bending modes. (**d**) VnNs show the characteristic weak absorbance band at 430nm.

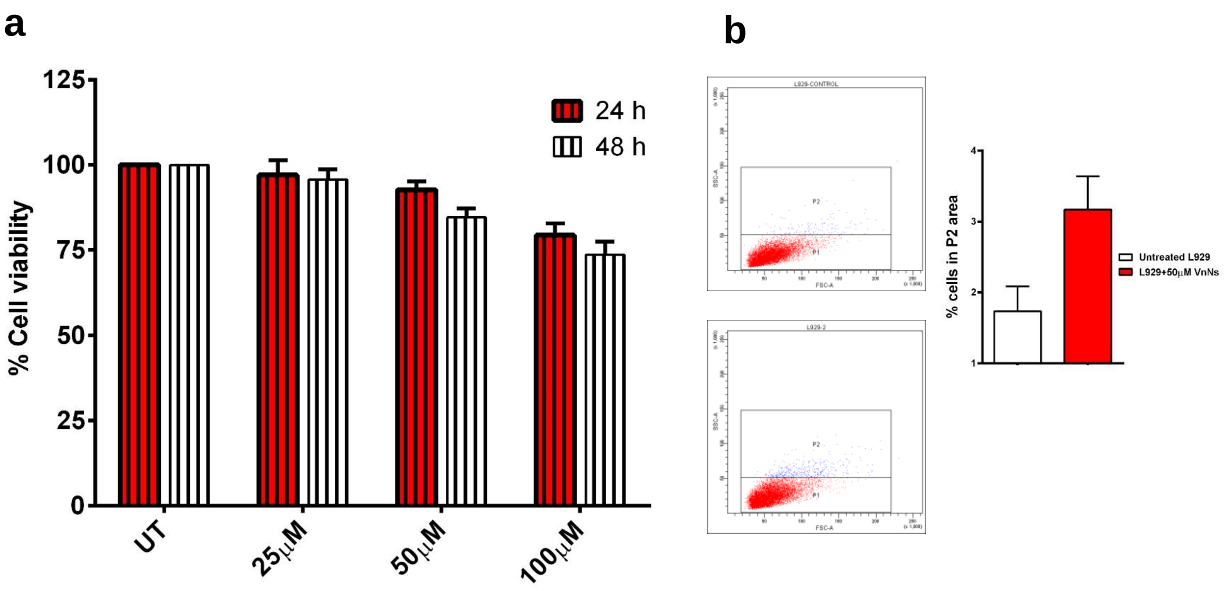


**Supplementary Figure 2. Cellular interaction studies on L929 cells.** (**a**) Demonstrate the percentage cell viability of L929 cells, post VnNp treatment towards 24 and 48h. (**b**) Changes in the side scatter pattern in L929 cells with VnNp treatment. The cells shown no significant increase in SSC (n=4 & non-significant P value).

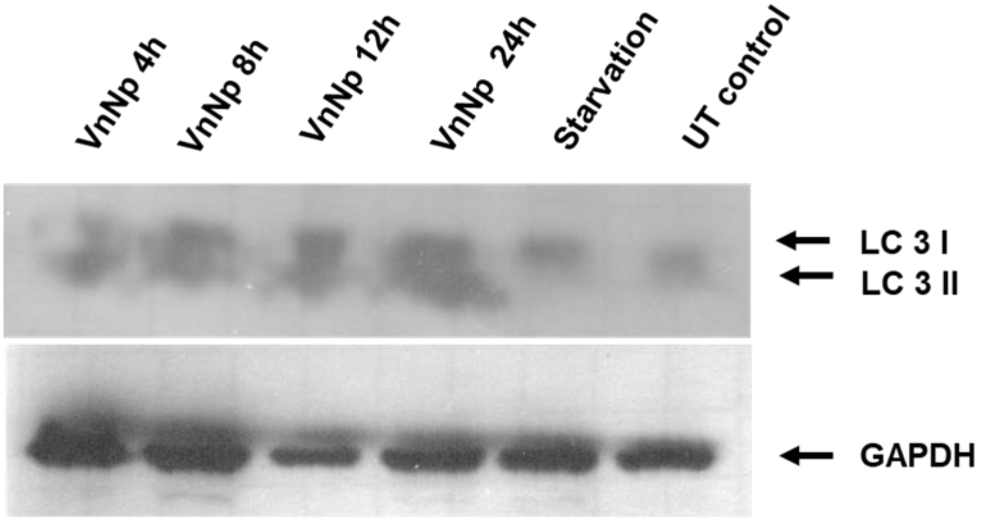


**Supplementary Figure 3. Western blot assay** elucidating the LC3 protein expression pattern in VnNp treated MDA-MB-231 cells at different time points. The image shows increased levels of both the LC 3 proteins in samples from all time point study as compared with the untreated control samples, which clearly indicate autophagy induction in cells at different time points until 24h. Lower panel on GAPDH depicts loading control.

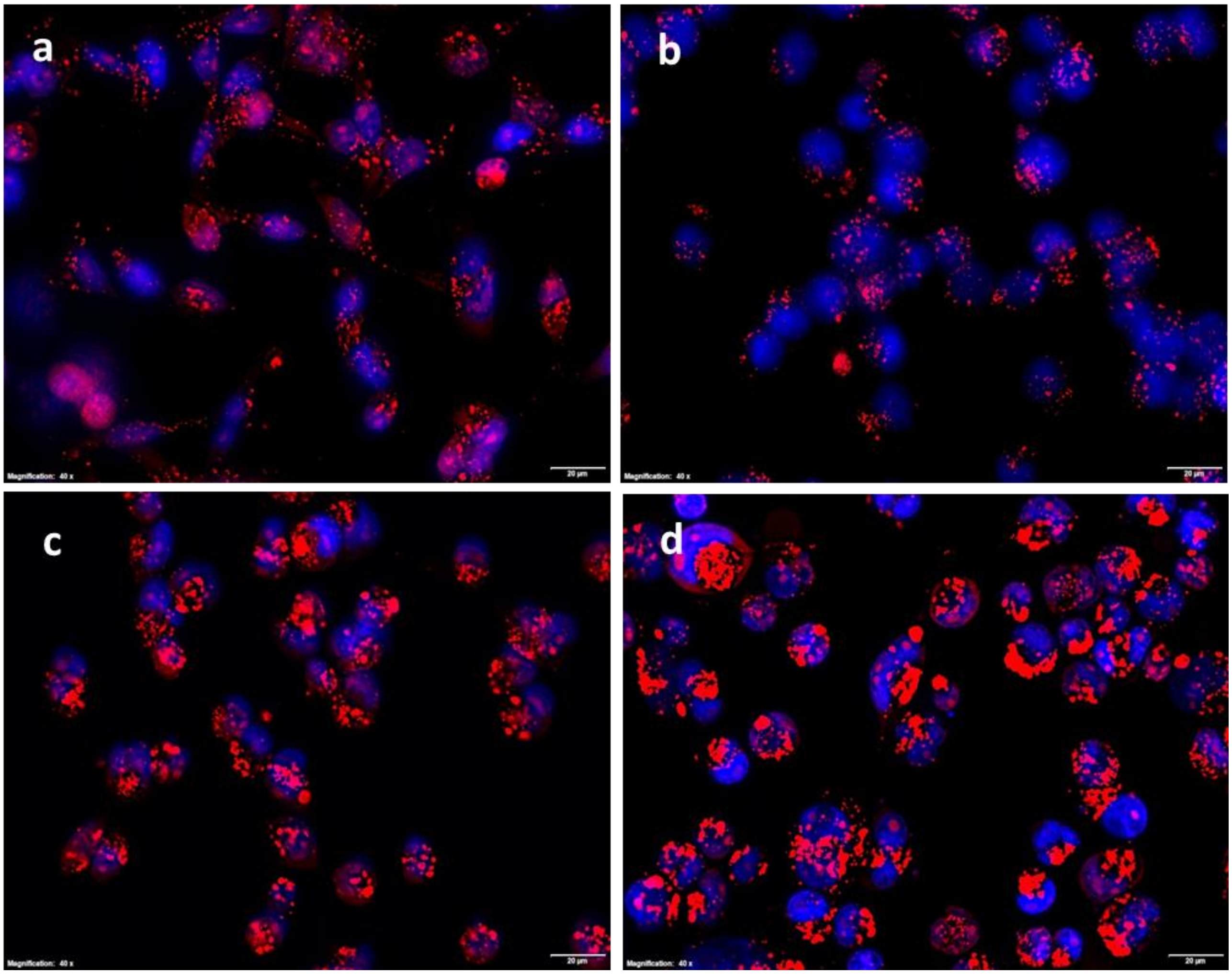


**Supplementary Figure 4. Lysosomal membrane integrity.** (**a**) Representative fluorescent micrographs of Acridine Orange stained 48h maintained untreated cells (**b**) cells treated with 50µM VnNp for 1h (**c**) 24h (**d**) 48h. Image shows acidic compartments with red sports giving a visual identification of VnNp induced increased lysosomal number, size and accumulation. n=4

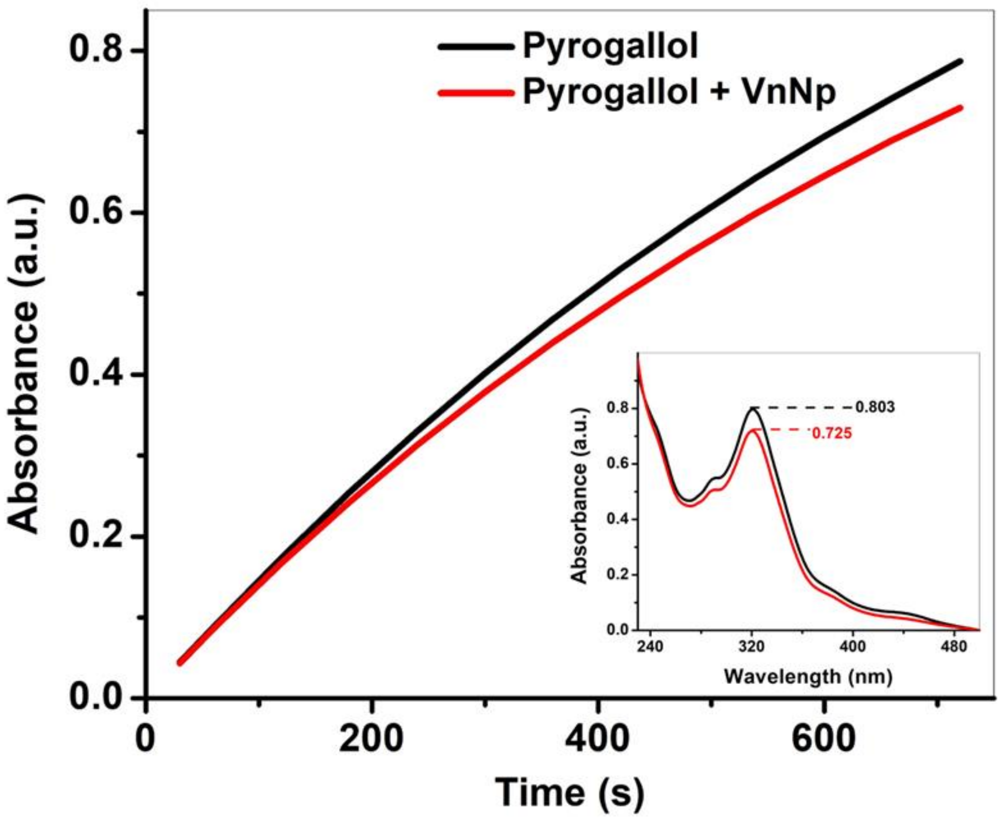


**Supplementary Figure 5**. **Kinetic study** on the inhibition of pyrogallol autoxidation in the presence of VnNp. Inset shows a decrease in absorbance pattern of purpurogallin, the product of pyrogallol auto-oxidation measure at 320nm.

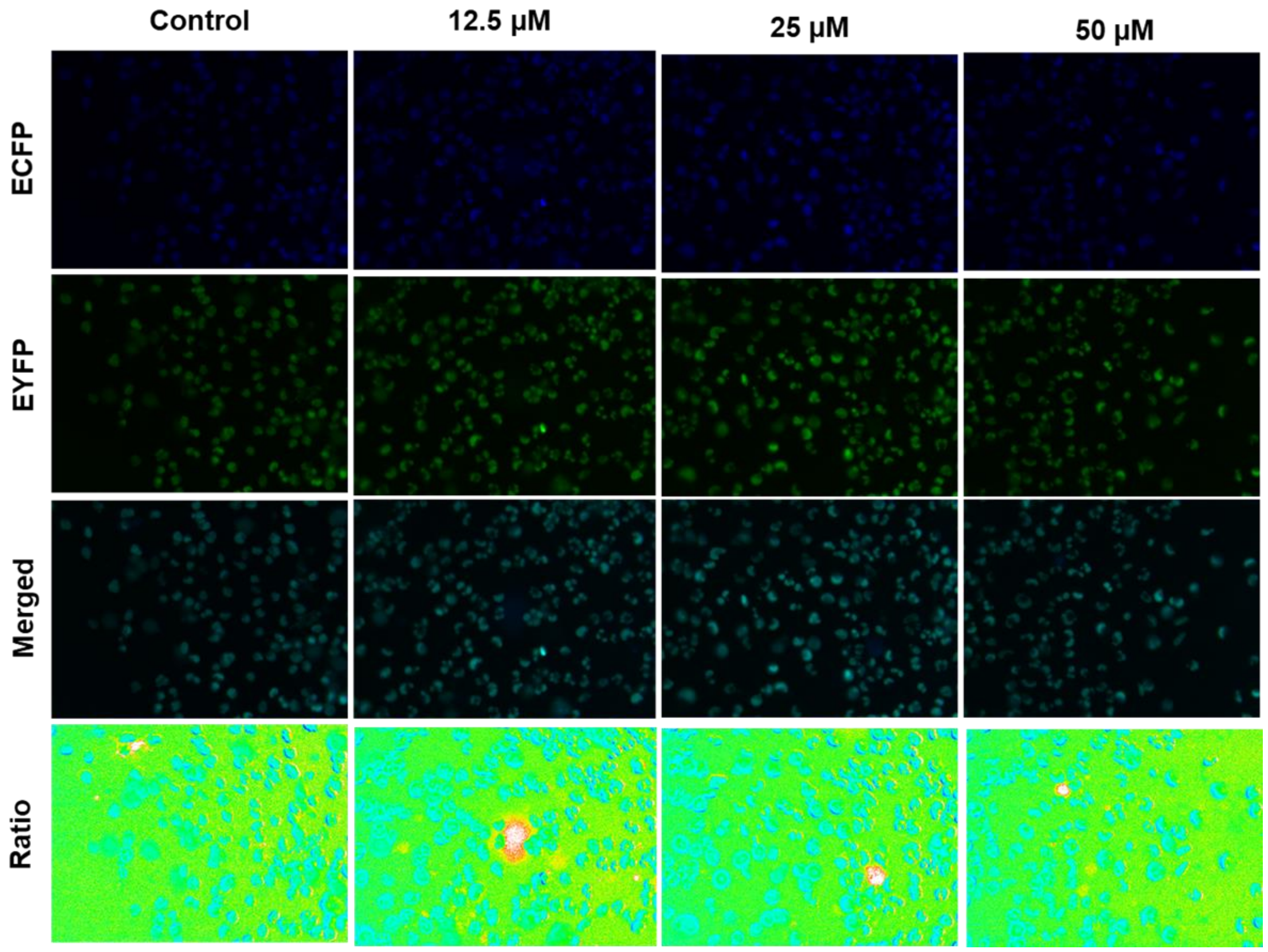


**Supplementary Figure 6. Caspase activity in SACT3 expression cells.** (**a**) Fluorescence micrographs showing the absence of FRET activity in cells treated with VnNp for 24h. None of the test groups indicated the onset of caspase activity.

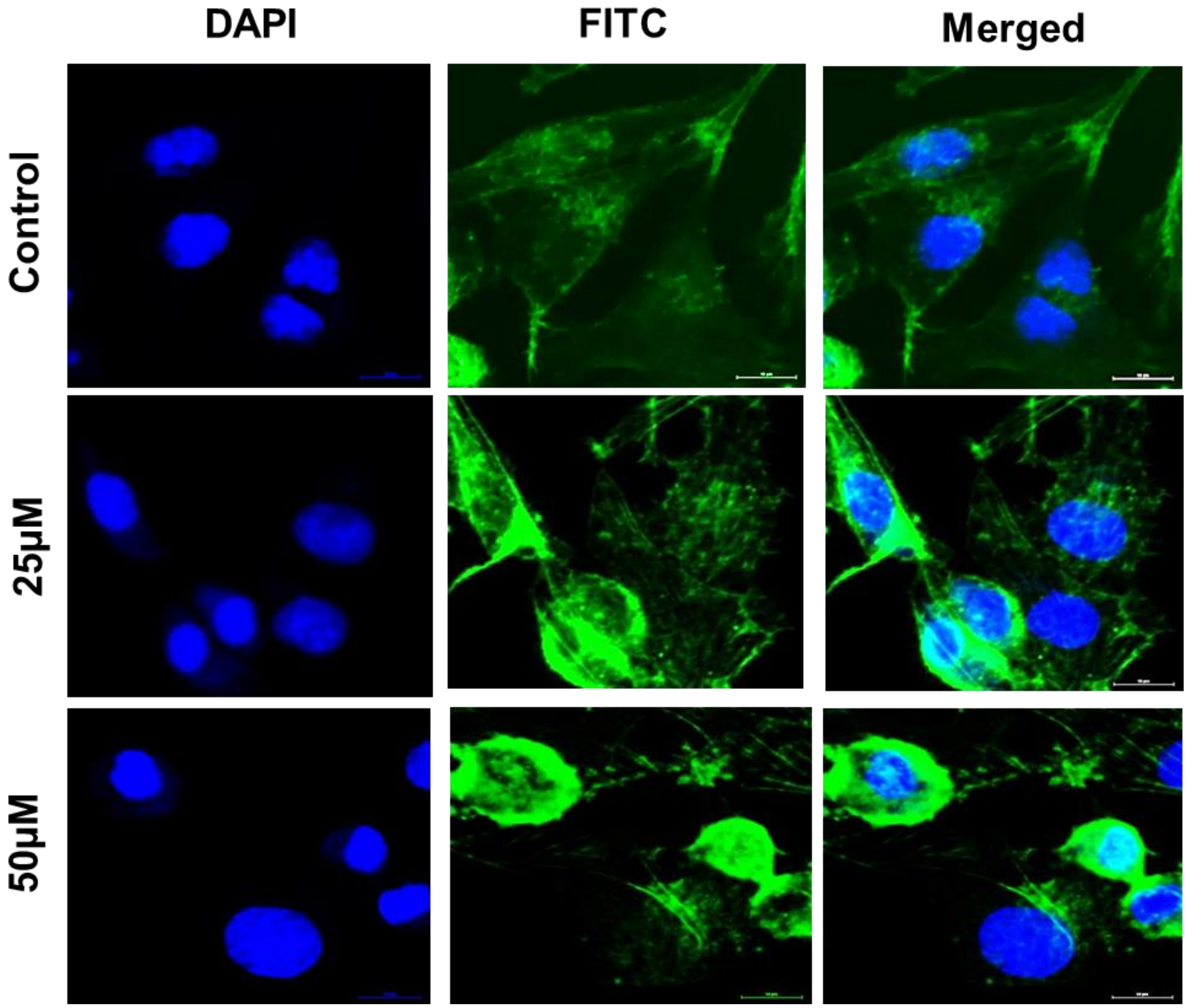


**Supplementary Figure 7. Morphology changes.** (**a**) Fluorescent micrographs depicting the well-structured actin organisation in untreated control cells. The 48h treated cells denote actin filament shrinkage with 25 and 50 µM treatment.

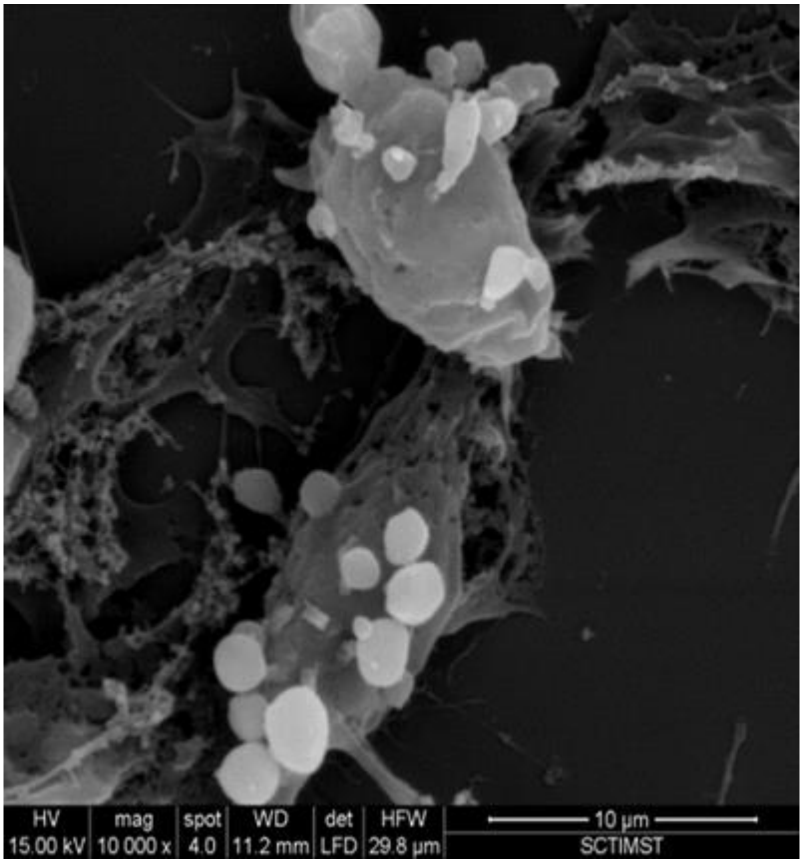


**Supplementary Figure 8. Membrane blebs in apoptosis.** Scanning electron micrograph depicting the characteristic cellular morphology in 50µM VnNp pretreated cells for 48h. Membrane blebbing is a characteristic to apoptosis.

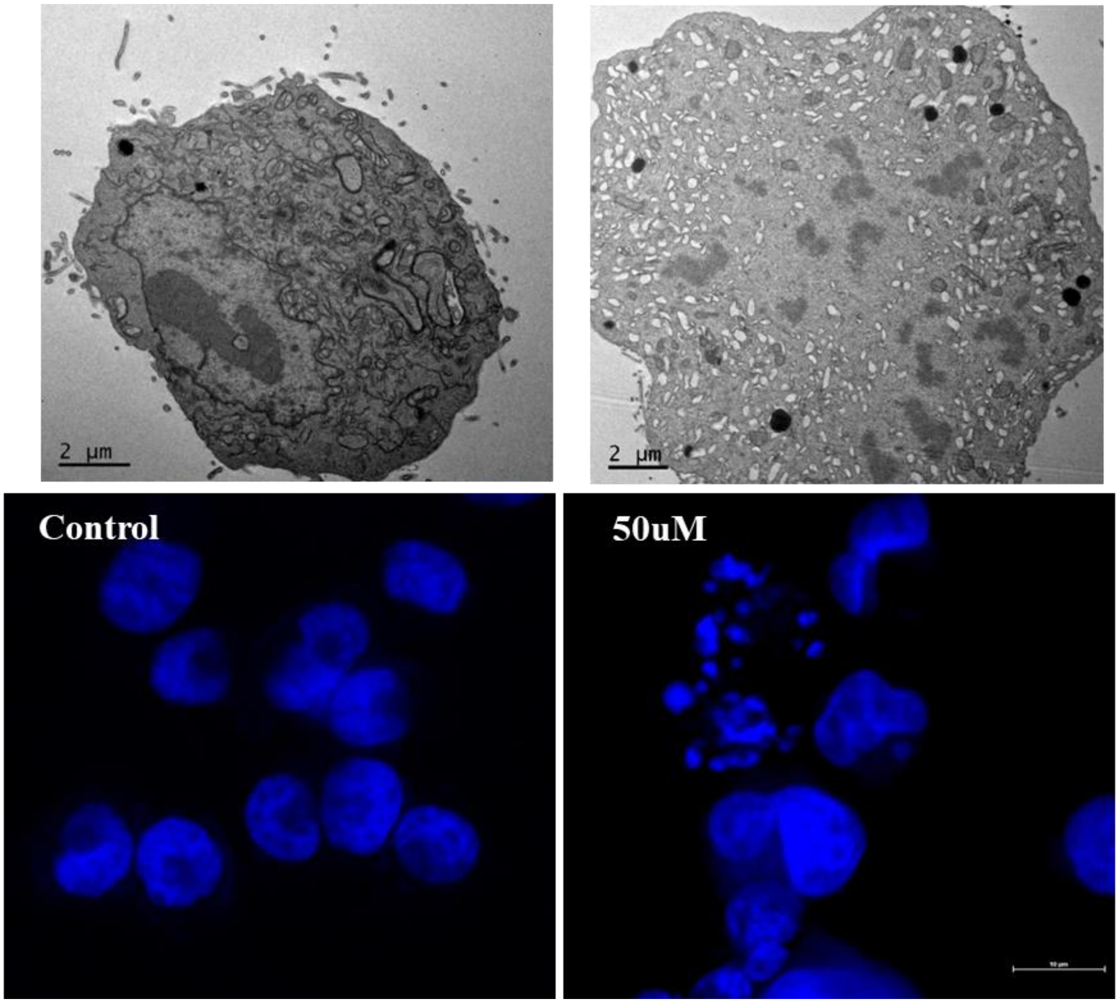


**Supplementary Figure 9. Nuclear morphology.** The TEM and fluorescence micrographs depict **c**hromatin condensation and breakdown of nuclear morphology, which are characteristic to apoptosis.

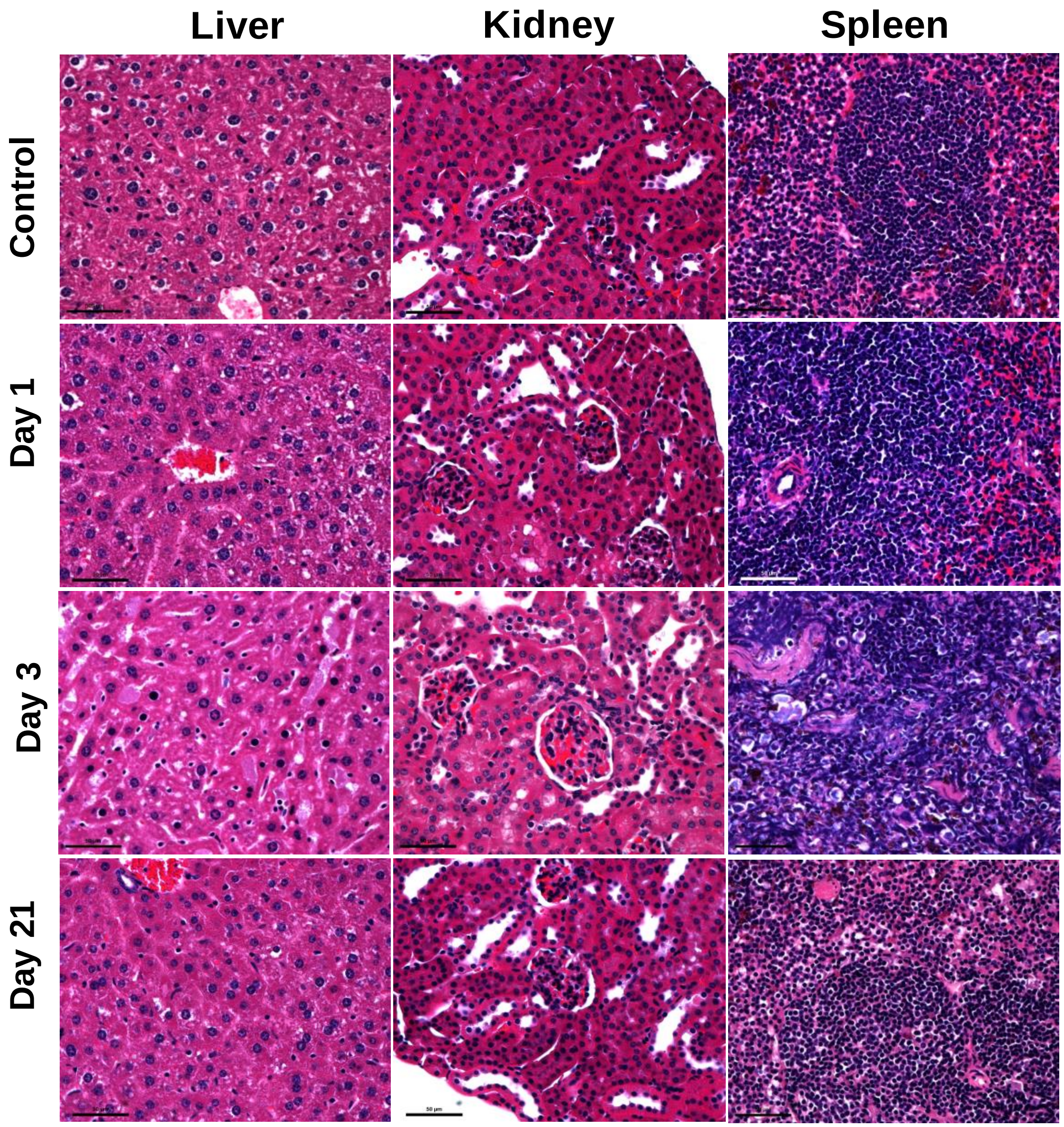


**Supplementary Figure 10. Histopathology of tissues.** Bright field micrographs depict tissue sections from liver, kidney and spleen from the 1mg/kg VnNp injected mice sacrificed at day-1, day-3 and day-21 post injection.

| Primer Name | Sequence |
|---|---|
| Mice | |
| m18SrNA_RT F | CGGAACTGAGGCCATGA |
| m18SrNA_RT R | CTTTCGCTCTGGTCCGTC |
| mCat_RT F | CGACCAGATGAAGCAGTGGAA |
| mCat_RT R | ACCCCGCGGTCATGATATTA |
| mSOD1_RT F | GAGACCTGGGCAATGTGACT |
| mSOD1_RT R | TTGTTTCTCATGGACCACCA |
| mSOD2_RT F | CTGGACAAACCTGAGCCCTA |
| mSOD2_RT R | GAACCTTGGACTCCCACAGA |
| mSOD3_RT F | AGTCCAGCTTCGACCTAGCA |
| mSOD3_RT R | CCATCCAGATCTCCAGCACT |
| Human | |
| hSOD1_RT F | GGTGTGGCCGATGTGTCTAT |
| hSOD1_RT R | CCTTTGCCCAAGTCATCTGC |
| hSOD3_RT F | AGGGACAGCCTGCGTTC |
| hSOD3_RT R | CAGGAACACAGTAGCGCCAG |
| hSOD2_RT F | GCTGCACCACAGCAAGCA |
| hSOD2_RT R | TCGGTGACGTTCAGGTTGTTC |

**Supplementary Table 1. List of primers.** Depict the list of RT-PCR primers used for the mRNA quantification on various tissue samples from Swiss albino mice and in MDA-MB-231 cells.
